# Comparative genomic analysis of *Bradyrhizobium* strains with natural variability in the efficiency of nitrogen fixation, competitiveness, and adaptation to stressful edaphoclimatic conditions

**DOI:** 10.1101/2024.01.10.574934

**Authors:** Milena Serenato Klepa, George Colin diCenzo, Mariangela Hungria

## Abstract

*Bradyrhizobium* is known for its ability to fix atmospheric nitrogen in symbiosis with agronomically important crops. This study focused on two groups of strains, each containing eight putative natural variants of *B. japonicum* SEMIA 586 (=CNPSo 17) or *B. diazoefficiens* SEMIA 566 (=CNPSo 10), previously used as commercial inoculants for soybean crops in Brazil. We aimed to detect genetic variations that might be related to biological nitrogen fixation, competitiveness for nodule occupancy, and adaptation to the stressful conditions of the Brazilian Cerrado soils. High-quality genome assemblies were produced for all strains and used for comparative genomic analyses. The core genome phylogeny revealed that strains of each group are closely related, confirmed by high average nucleotide identity (ANI) values. However, variants accumulated divergences resulting from horizontal gene transfer (HGT), genomic rearrangements, and nucleotide polymorphisms. The *B. japonicum* group presented a larger pangenome and a higher number of nucleotide polymorphisms than the *B. diazoefficiens* group, probably due to its longer adaptation time to the Cerrado soil. Interestingly, five strains of the *B. japonicum* group carry two plasmids. The genetic variability found in both groups is discussed in light of the observed differences in their nitrogen fixation capacity, competitiveness for nodule occupancy, and environmental adaptation.

**SIGNIFICANCE:** The two main reference strains for soybean inoculation in Brazil, *B. japonicum* CPAC 15 (=SEMIA 5079) and *B. diazoefficiens* CPAC 7 (=SEMIA 5080), have been considered highly competitive and highly efficient in nitrogen fixation, respectively. In this study, we obtained and analyzed the genomes of the parental and variant strains. We detected two plasmids in five strains and several genetic differences that might be related to adaptation to the stressful conditions of the soils of the Brazilian Cerrado biome. We also detected genetic variations in specific regions that may impact symbiotic nitrogen fixation. Our analysis contributes to new insights into evolution of *Bradyrhizobium*, and some of the identified differences may be applied as genetic markers to assist strain selection programs.

## INTRODUCTION

Agriculture faces global challenges to feed the world’s growing population in the coming years (United Nations, 2017). Major concerns associated with the need for increased crop production relate to the degradation of natural ecosystems and the emission of greenhouse gases (Tilman et al., 2011). Technologies based on plant growth-promoting bacteria have high potential to support the development of more sustainable agricultural practices. Biological nitrogen fixation (BNF) refers to the reduction of the atmospheric nitrogen (N_2_) into assimilable forms by a diverse group of prokaryotic organisms. When performed in association with plants, the fixed nitrogen (N) may be used by the plants to help meet their N demand. In terrestrial ecosystems, the primary source of BNF is the N-fixing endosymbiosis between legume plants and bacteria collectively known as rhizobia, with the global contribution of symbiotic N fixation estimated at around 35 million tons of N fixed in 2018 (Herridge et al., 2022).

Brazil is a leader in the application of elite *Bradyrhizobium* strains as inoculants in soybean (*Glycine max* (L.) Merrill) fields. In the 2022/2023 crop season, around 44 million hectares were cropped with soybean, with 110 million doses of inoculants applied. Considering the methodology of Telles et al. (2023), economic savings due to the replacement of N-fertilizers with BNF in Brazilian soybean crops was US$27.4 billion in 2022/2023, in addition to mitigating 236 million equivalents of CO_2_.

The success of BNF in Brazil results from a long-term strain selection program, which started by looking for variants of exotic introduced *Bradyrhizobium* strains adapted to tropical conditions and Brazilian soybean genotypes (Hungria and Mendes, 2015). Currently, the elite strains *B. japonicum* CPAC 15 (=SEMIA 5079) and *B. diazoefficiens* CPAC 7 (= SEMIA 5080), two such natural variants, compose most of the soybean commercial inoculants. Numerous other natural variants of past inoculant strains have been also isolated from Brazilian soils and today are recommended for other legumes or grasses (Hungria et al., 2000; Hungria et al., 2010; Hungria and Mendes, 2015). The variants differ in their N-fixation capabilities and competitiveness for legume nodule occupancy, although in most cases, the genetic variations responsible for the phenotypic differences are not yet known.

Species of the genus *Bradyrhizobium* display broad variation in lifestyles and legume host range, presumably influencing the group’s genetic organization (Ormeño-Orrillo and Martínez-Romero, 2019). Conceptually, pangenomes represent the entire set of genes of a phylogenetically related group of strains, fractioned into core and dispensable genomes (Tettelin et al., 2008). The core genome consists of genes shared by all strains of the group and usually codes essential functions. In contrast, the dispensable genome refers to genes present in a subset of strains (the accessory genome) or only in individual strains (the unique genome), and it is composed of genes that may provide additional functions for those strains and often relate to environmental adaptation (Vernikos et al., 2015). Analyses of bacterial pangenomes may elucidate genomic variability even between closely related strains within a species, and reveal information about horizontal gene transfer (HGT) and evolution (Medini et al., 2005).

Events of HGT, recombination, and mutations are some of the main drivers of bacterial evolution (Arnold et al., 2022). HGT refers to the transfer of DNA segments from one organism to another. Mobile genetic elements such as plasmids and integrative conjugative elements (ICE) may contribute to the fitness of recipient bacteria by conferring selective advantages, including legume symbiosis, antibiotic resistance, and pathogenicity (Juhas et al., 2008; Weisberg et al., 2022). Concerning the genomes of rhizobia, symbiotic plasmids (i.e., plasmids encoding the determinants of legume symbiosis) are most prevalent in the genera *Sinorhizobium* and *Rhizobium*, whereas the genera *Bradyrhizobium* and *Mesorhizobium* usually carry their symbiotic genes in genomic islands integrated into the chromosomes, known as symbiosis islands (SI) (Wang et al., 2019; Weisberg et al., 2022). Genomic islands are commonly inserted at tRNA genes and may present a lower GC mol % in comparison to the remaining chromosome, in addition to a high number of genes encoding hypothetical proteins, plasmid conjugation, integrases, insertion sequences (IS), and transposases (Juhas et al., 2008). This movement of DNA segments may prompt genome recombination, including inversions, translocations, duplications, nucleotide polymorphisms, and indels, resulting in genetic variability even between closely related strains (Hughes, 2000).

In this study, we evaluated the genetic variability of two groups of closely related natural variants of *B. japonicum* and *B. diazoefficiens* adapted to Brazilian Cerrado soils – the main cropping area in the country – for longer or shorter periods, respectively. Each group included the parental strain previously used as a commercial inoculant in Brazil, one natural variant strain currently used in commercial inoculants and treated as the reference genome, and seven other putative natural variants. The *B. japonicum* strains were previously shown to vary in their competitiveness for nodule occupancy, while the *B. diazoefficiencs* strains differ in their capacity for BNF (**Table 1)**. Previously, a chromosome-level genome sequence existed only for strain CPAC 15 (reference strain for the *B. japonicum* group) (Siqueira et al., 2014). Here, we report complete or high-quality draft genomes for the remaining 17 strains. Subsequent comparative genomic analyses allowed for the identification of genetic differences that may explain the observed differences in their capacity for BNF capacity and competitiveness for nodule occupancy.

**Table 1.** Competitiveness and BNF capacity comparison between the parental strain (P) and the variant strains compared to the reference variant (R). Data from the CPAC 15 group was retrieved from Hungria et al. (1998), while data for the CPAC 7 group was retrieved from Santos et al. (1999).

|  | Strains | Synonym | Competitiveness | BFN capacity |
| --- | --- | --- | --- | --- |
| *Bradyrhizobium japonicum*<br>**CPAC 15** (=SEMIA 5079,<br>CNPSo 7, DF 24) (R) | CNPSo 17 (P) | SEMIA 566, BR 40 | = | < |
|  | CNPSo 22 | S340 | > | < |
|  | CNPSo 23 | S370 | > | > |
|  | CNPSo 24 | S372 | > | > |
|  | CNPSo 29 | S478 | > | > |
|  | CNPSo 31 | S490 | > | < |
|  | CNPSo 34 | S516 | > | < |
|  | CNPSo 38 | S204 | = | < |
| *Bradyrhizobium diazoefficiens*<br>**CPAC 7** (= SEMIA 5080,<br>CNPSo 6) (R) | CNPSo 10 (P) | SEMIA 586, CB 1809 | < | = |
|  | CNPSo 104 | CPAC 390 | > | > |
|  | CNPSo 105 | CPAC 392 | < | < |
|  | CNPSo 106 | CPAC 393 | = | = |
|  | CNPSo 107 | CPAC 394 | = | = |
|  | CNPSo 108 | CPAC 402 | = | > |
|  | CNPSo 109 | CPAC 403 | = | = |
|  | CNPSo 110 | CPAC 404 | < | = |

## RESULTS

### Whole genome sequencing statistics

Hybrid genome assembly using Oxford Nanopore Technologies (ONT) and Illumina reads from the *B. japonicum* strains (excluding CPAC 15, for which the published genome was used) resulted in seven finished genomes and one strain with the chromosome split into two contigs (**Table 2**). The genomes ranged from 9,584,431 bp to 10,367,856 bp, which is comparable to the published genome of the reference strain CPAC 15, estimated at 9,583,027 bp (Siqueira et al., 2014), and of the type strain USDA 6^T^ with 9,207,384 bp (Kaneko et al., 2011). The total number of coding genes ranged from 8,758 to 9,472. The GC content (mol %) ranged from 63.34% to 63.55%.

**Table 2.** Genome assembly statistics of the parental strain (P), the reference variant for the genome comparison (R), and other variant strains of the *B. japonicum* and *B. diazoefficiens* groups.

| Strain | GenBank accession number | Size (bp) | Contigs | Number of plasmids | Coverage (-fold) | CDS with protein | Pseudogenes | ANI against CPAC 15 |
| --- | --- | --- | --- | --- | --- | --- | --- | --- |
| *B. japonicum* group |  |  |  |  |  |  |  |  |
| CNPSo 17 (P) | CP136588 | 9,609,914 | 1 | 0 | 102 | 8,796 | 301 | 99.9872 |
| CNPSo 22 | CP141637-CP141639 | 10,295,385 | 3 | 2 | 206 | 9,335 | 439 | 99.9391 |
| CNPSo 23 | CP139647 | 9,586,320 | 1 | 0 | 146 | 8,758 | 310 | 99.9961 |
| CNPSo 24 | CP138298 | 9,584,431 | 1 | 0 | 76 | 8,765 | 299 | 99.9966 |
| CNPSo 29 | CP138299-CP138301 | 10,284,125 | 3 | 2 | 63 | 9,303 | 447 | 99.9359 |
| CNPSo 31 | CP138302-CP138304 | 10,280,370 | 3 | 2 | 225 | 9,305 | 434 | 99.9392 |
| CNPSo 34 | CP139648-CP139650 | 10,357,443 | 3 | 2 | 144 | 9,467 | 433 | 99.9266 |
| CNPSo 38 | JAXIOH000000000 | 10,367,856 | 4 | 2 | 94 | 9,472 | 447 | 99.922 |
| CPAC 15 (R) | CP007569 | 9,583,027 | 1 | 0 | 20 | 8,653 | 336 | 100 |
| Strain | GenBank accession number | Size (bp) | Contigs | Number of plasmids | Coverage (-fold) | CDS with protein | Pseudogenes | ANI against CPAC 7 |
| *B. diazoefficiens* group |  |  |  |  |  |  |  |  |
| CNPSo 10 (P) | CP139628 | 9,140,459 | 1 | 0 | 83 | 8,230 | 280 | 99.9986 |
| CNPSo 104 | CP139629 | 9,140,039 | 1 | 0 | 168 | 8,231 | 285 | 99.9978 |
| CNPSo 105 | CP139630 | 9,140,270 | 1 | 0 | 95 | 8,236 | 269 | 99.9988 |
| CNPSo 106 | CP139631 | 9,120,098 | 1 | 0 | 60 | 8,220 | 270 | 99.9971 |
| CNPSo 107 | CP139632 | 9,138,100 | 1 | 0 | 37 | 8,224 | 279 | 99.9983 |
| CNPSo 108 | CP139633 | 9,138,003 | 1 | 0 | 41 | 8,235 | 270 | 99.9989 |
| CNPSo 109 | CP139634 | 9,138,085 | 1 | 0 | 109 | 8,244 | 264 | 99.9985 |
| CNPSo 110 | CP139635 | 9,138,906 | 1 | 0 | 109 | 8,230 | 275 | 99.9988 |
| CPAC 7 (R) | CP139636 | 9,138,870 | 1 | 0 | 66 | 8,235 | 275 | 100 |

The genome assemblies from the *B. diazoefficiens* group resulted in finished genomes for all nine strains (**Table 2**). The genome sizes were smaller than those of the *B. japonicum* group, ranging from 9,120,098 bp to 9,138,003 bp, which is very close to the genome size of the type strain USDA 110^T^ (9,105,828 bp) (Kaneko et al., 2002), and of the original draft genome of reference strain CPAC 7, estimated at 9,138,870 bp (Siqueira et al., 2014). The total number of coding genes ranged from 8,220 to 8,244, and the GC content (mol %) from 63.96% to 63.97%.

We computed the average nucleotide identity (ANI) of each parental and variant against the reference strain of each group (CPAC 15 and CPAC 7). The strains of both groups shared high ANI values, confirming that they are closely related. The *B. japonicum* strains shared values equal to or higher than 99.82 % with CPAC 15, and all strains of the *B. diazoefficiens* group shared ∼99.99 % with CPAC 7. ANI comparisons against the species type strains, *B. japonicum* USDA 6^T^ and *B. diazoefficiens* USDA 110^T^, confirmed the species designations of all strains were accurate (**Table 2**).

### Core genome phylogeny and genome synteny

As expected, the *B. japonicum* and *B. diazoefficiens* groups were separated into two clades with high bootstrap support in a core-genome phylogeny **(Figure 1)**. The type strains *B. japonicum* USDA 6^T^ and *B. diazoefficiens* USDA 110^T^ were included in the phylogeny and presented a basal position in the respective clade of each species. Since we are working with closely related strains in each group, the core genome phylogeny showed low differentiation between the strains of both groups. Interestingly, the *B. japonicum* group could be sub-divided into two well-supported monophyletic groups. The five variants of the *B. japonicum* group with a genome size > 10.2 Mb (compared to ∼ 9.6 Mb for the other *B. japonicum* strains) formed one monophyletic group, suggesting an ancestral genome enlargement at the base of this clade, while the other four strains (including CPAP 15) formed a second monophyletic group **(Figure 1)**.

**Figure 1.**
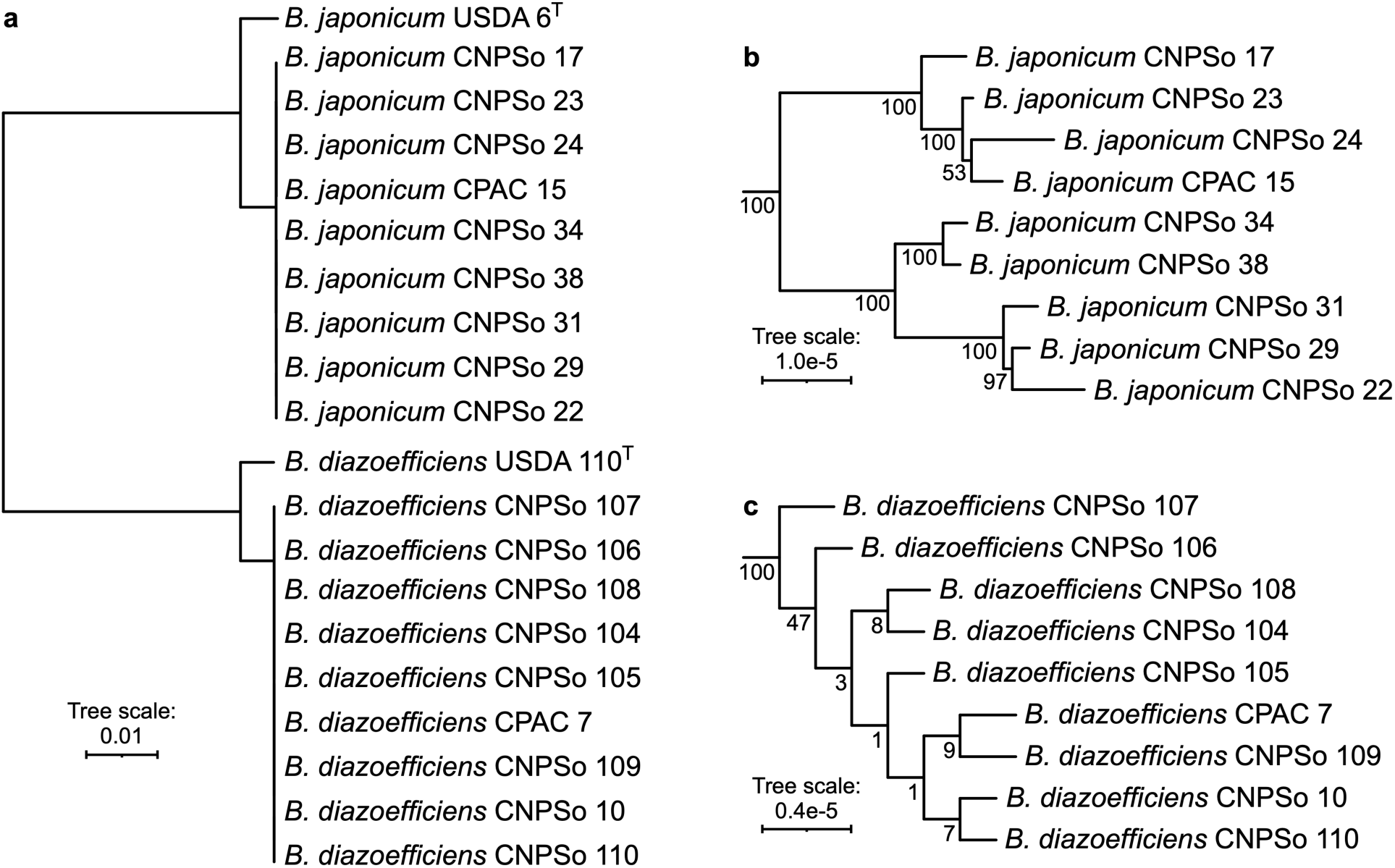
Unrooted phylogeny of reference, parental, and variant strains of the *B. japonicum* and *B. diazoefficiens* groups. (**a**) A maximum likelihood phylogeny of all strains was prepared from a concatenated alignment of 2,689 core genes. The scale represents the mean number of nucleotide substitutions per site. Sub-trees of the (**a**) the *B. japonicum* and (**b**) the *B. diazoefficiens* groups are shown with a different branch length scale to better display the within-group relationships. Numbers at the nodes indicate the bootstrap values, based on 1,000 bootstrap replicates.

The genomes of the *B. japonicum* group are highly syntenic; however, it is possible to observe rearrangements, including inversions, transpositions, and additions/deletions **(Figure 2a).** By comparing the genome of the *B. japonicum* reference strain CPAC 15 and its parental CNPSo 17, we observed a possible inversion of a large segment (around 3,500,000 bp). The strains CNPSo 23, CNPSo 24, CNPSo 29, CNPSo 31, CNPSo 34, and CNPSo 38 are highly syntenic with CNPSo 17, indicating conservation of the gene order. On the other hand, strain CNPSo 22, which is polyphyletic with CPAC 15, appears to have the same inversion present in CPAC 15. However, we note that CPAC 15 and CNPSo 22 are the only genomes not assembled using Flye, and thus we cannot rule out that the putative inversion instead reflects assembly errors. Other small inversions were detected, as well as translocations and deletions; however, they are unlikely to impact the symbiosis islands **(Figure 2a)**. We found a higher number of IS and transposases in the five larger genomes. *B. japonicum* strains CNPSo 22, CNPSo 29, CNPSo 31, CNPSo 34, and CNPSo 38 (genome size > 10.2 Mb) have around 121 ISs and 29 transposase genes, whereas the strains CPAC 15, CNPSo 17, CNPSo 23, and CNPSo 24 (genome size ∼ 9.6 Mb) contain around 68 and 10, respectively.

**Figure 2.**
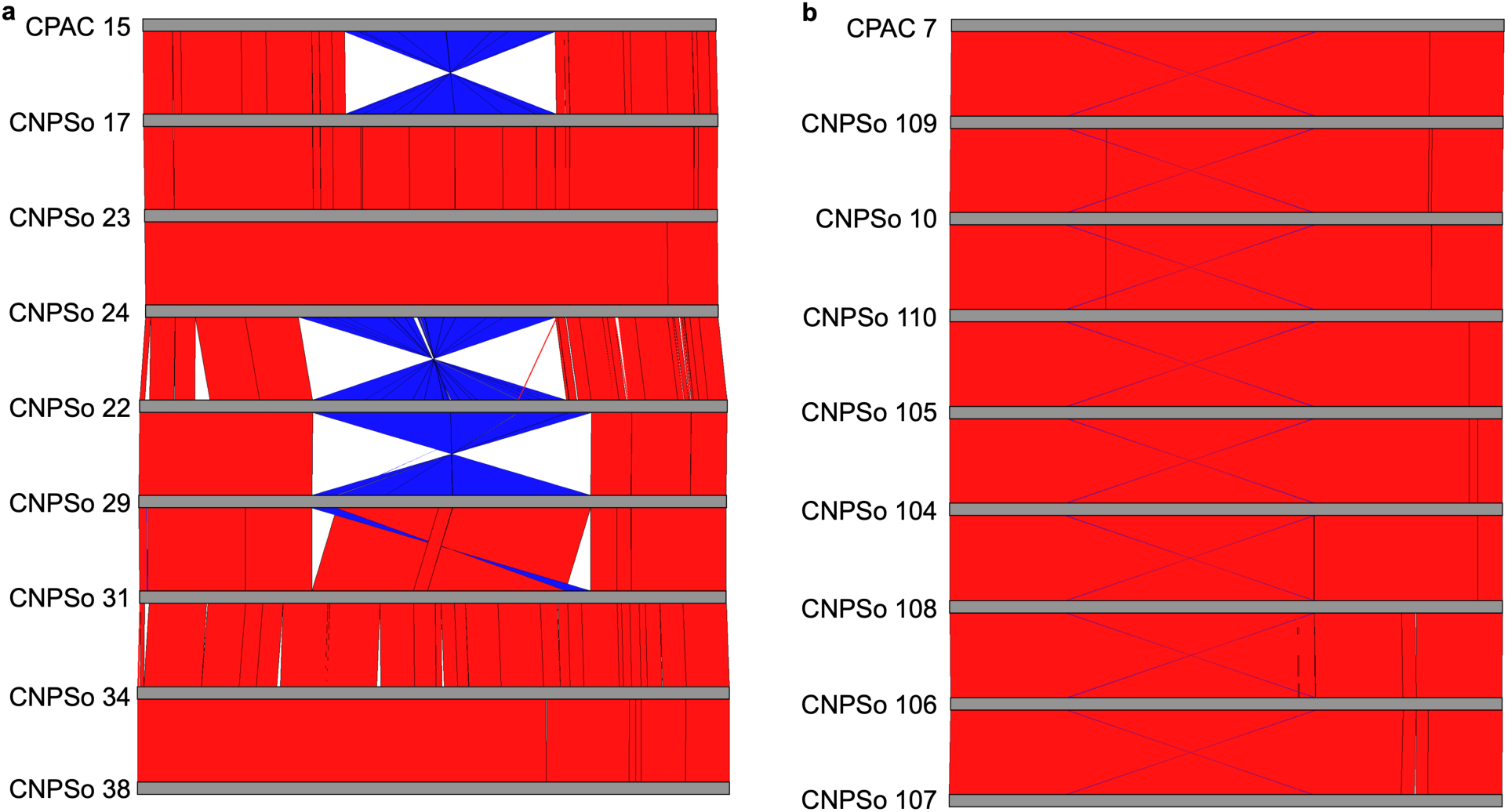
Genome-wide synteny analysis of the (**a**) *B. japonicum* and (**b**) *B. diazoefficiens* parental strains (CNPSo 17 and CNPSo 10), reference strains (CPAC 15 and CPAC 7), and other variants. Pairwise genome alignments were performed, and homologous regions between pairs of genomes are connected by red lines (if in the same orientation) or blue lines (if in the inverse orientation).

The genomes of the *B. diazoefficiens* group are highly syntenic **(Figure 2b)**, and unlike the *B. japonicum* group, displayed no relevant rearrangements. The strains of the *B. diazoefficiens* group contained 69 ISs and nine transposase genes.

### Plasmid content

We identified two sets of orthologous plasmids in the five *B. japonicum* strains with larger genomes (strains CNPSo 22, CNPSo 29, CNPSo 31, CNPSo 34, and CNPSo 38), which were confirmed as plasmids by the presence of a *repABC* operon encoding proteins responsible for plasmid replication and partitioning (Cevallos et al., 2008). Plasmid “a” presented high similarity among the strains, with 98 to 99.9% of identity and sizes ranging from 160,017 bp to 170,161 bp. Plasmid “b” shared 96.6 to 99.9% identities among the strains and sizes ranging from 283,210 bp and 291,690 bp. Interestingly, the plasmids did not account for the full difference in the genome sizes of these five *B. japonicum* strains compared to the four with smaller genomes (CPAC 15, CNPSo 17, CNPSo 23, and CNPSo 24), indicating that this lineage also experienced an ancestral chromosome enlargement. No plasmids were found in the strains of the *B. diazoefficiens* group.

A BLASTn search showed that a segment (∼ 49,500 bp) of plasmid “a” of all variants shares >99.95% identity to the 210 kb plasmid pNK6b (GenBank accession AP014686.1) of *B. diazoefficiens* NK6 (Iida et al., 2015). The number of genes on the “a” plasmids ranged from 123 to 132, and encoded a type III secretion system (T3SS), which is known to participate in host cell infections, including symbiotic interactions (Teulet et al., 2022). The T3SS cluster of plasmid “a” is composed of the secretion and cellular translocation (*sct*) genes (*sctNVJQRSTU*) in all strains except for CNPSo 29, which contained a putative deletion of eight genes caused by an IS4 family transposase, including part of T3SS apparatus. Interestingly, plasmid “a” also harbors genes related to stress responses, including genes coding for toxin-antitoxin systems (TA systems) (MazE/MazF, Phd/YefM), a secretion protein (HlyD family efflux transporter periplasmic adaptor subunit; lost in CNPSo 38), a heat-shock protein (molecular chaperone HtpG), and a trehalose-6-phosphate synthase. A LysR substrate-binding domain-containing protein related to nitrogen metabolism was also identified. Moreover, plasmid “a” contained the gene clusters *traADGFHM* and *trbBCDEFGHIJKL* related to conjugation.

The entire sequence of plasmid “b” shared around 99.99% identity with the 290 kb plasmid pN03G-2 of *B. japonicum* pN03G-2 (GenBank accession CP126012), suggesting common ancestry. The number of genes on the “b” plasmids ranged from 212 to 219, and the annotation revealed genes encoding proteins related to stress responses, such as TA systems (VapB/VapC, Phd/YefM, AbiEii/AbiGii), efflux transporters (HlyD and HlyB families), a cold-shock protein, heat-shock proteins (chaperonin GroEL, co-chaperone GroES), and an O-antigen gene cluster. Plasmid “b” also carries several genes encoding proteins involved in the synthesis, transport, metabolism, and regulation of amino acids, nucleic acids, carbohydrates, and lipids. Even though no T3SS genes were detected, two T3SS effector proteins (C48 family peptidase and E3 ubiquitin--protein ligase) were identified. The conjugation apparatus of plasmid “b” is encoded by *traACDFGM* and *trbBCDEJLFGIH*. More information about the gene contents of the plasmids is presented in **Table S1.**

### Symbiosis islands

The symbiosis islands (SIs) A and B of both groups were detected according to Weisberg et al. (2022), whereas the SI C was identified as the region proposed by Kaneko et al. (2011). In addition, the automatic annotation of SI A was manually curated according to previous studies (Kaneko et al., 2002; Teulet et al., 2020; Weisberg et al., 2022). SI A of the *B. japonicum* and *B. diazoefficiens* groups were bordered by tRNA-valine and a recombinase gene. Within each group of strains, the SIs are highly syntenic **(Figure S1; Figure S2)**, although some differences in gene content are evident.

The average size of SI A in the *B. japonicum* genomes is ∼690,000 bp and can be found between the chromosomal positions 7,581,129 and 8,615,546 depending on the strain. SI A contains between 584 to 615 genes. SI A of all *B. japonicum* strains carried the genes classically important for nodulation and nitrogen fixation, including: *nodD2D1ABCSUIJZ*, *noeEIL, nolAIKNOY, nolUV*, the regulatory system *nodV/nodW*, the nitrogen fixation genes *fixABCKRWX* and *nifABDEHKNQSTWXZ*, T3SS genes (*sct*, also referred to as ‘*Rhizobium*-conserved’ genes, *rhc*) *rhcC1C2DJNQRSTUV,* and genes encoding T3SS effector proteins known as nodulation outer proteins (*nop*), *nopAALARBWE1HLMP1P2P3.* In addition to these symbiosis genes, between 155 and 222 hypothetical genes, 75 IS from diverse families of transposases, and 100 pseudogenes were annotated. Interestingly, the monophyletic group of strains carrying plasmids had a smaller SI A (28 fewer genes on average) than those without plasmids **(Figure S1a)**; none of the known symbiosis genes were absent from SI A in these strains. SI B of the *B. japonicum* group, located between chromosomal positions 1,465,741 and 1,809,591 depending on the strain, is entirely syntenic among the strains (data not shown) and contains 4,160 bp with seven genes; an integrase gene and *ybgC* served as the borders of this SI, with five intervening hypothetical genes. With an average size of 203,000 bp, SI C was also highly syntenic and smaller in the strains carrying plasmids (18 fewer genes on average) **(Figure S1b)**. SI C can be found between chromosomal positions 8,807,943 and 9,339,632 depending on the strain. The region is bordered by genes coding for a tyrosine-type recombinase/integrase and a 5’-methylthioadenosine/S-adenosylhomocysteine, and includes ∼225 genes, with 59 to 65 hypothetical genes, 36 pseudogenes, and 24 IS of various transposase families. The average GC mol content of SI A, B, and C of the *B. japonicum* group was 60.58%, compared to 63.34% to 63.55% for the genome-wide average.

SI A of the *B. diazoefficiens* group is slightly smaller than in the *B. japonicum* group, with an average size of 671,500 bp. It is located between the chromosomal positions 7,426,338 and 8,082,430. The average number of genes within SI A is 570, including 155 hypothetical genes, 85 IS from diverse transposase families, and 97 pseudogenes. This region in *B. diazoefficiens* is highly syntenic, except for a ∼2 kb deletion in the genome of CNPSo 106 **(Figure S2a)**. The nodulation, nitrogen fixation, and T3SS genes are preserved as in the *B. japonicum* group. SI B, which is conserved across strains (data not shown), is 15,546 bp in length and is located between chromosomal positions 1,661,452 and 1,677,009. This region contains 15 genes, including nine hypothetical genes, one pseudogene, and one IS5 family transposase. SI C of the *B. diazoefficiens* group is found between the chromosomal positions 620,318 and 776,578 in the genomes and is ∼156,300 bp in length. The average number of genes is 156, with 49 hypothetical genes, 31 pseudogenes, and 16 IS from different transposase families. This region also displays high synteny across the strains **(Figure S2b)**. The average GC mol content of SI A, B, and C of the *B. diazoefficiens* group was 59.96%, compared to 63.96% to 63.97% for the genome-wide average.

### Pangenome analysis

A pangenome analysis was performed to understand the genomic variability of the strains. The pangenome of the *B. japonicum* group comprises 10,550 genes, 8,787 of which belonging to the core genome, 1,559 to the accessory genome, and 204 unique genes. Interestingly, 748 genes were shared only between the monophyletic group of strains carrying the plasmids; in contrast, only 187 genes were shared only between the strains without a plasmid **(Figure 3a)**. In addition, strain CNPSo 22 carries the largest number of unique genes (44), followed by CNPSo 17 (31), and then CNPSo 29 and CNPSo 38 (each with 27). Of the dispensable genome fraction, 148 genes (125 accessory genes and 23 unique genes) belong to SI A and thus are the most likely to contribute to the differences in BNF capacity and competitiveness of these strains **(Table S2)**.

**Figure 3.**
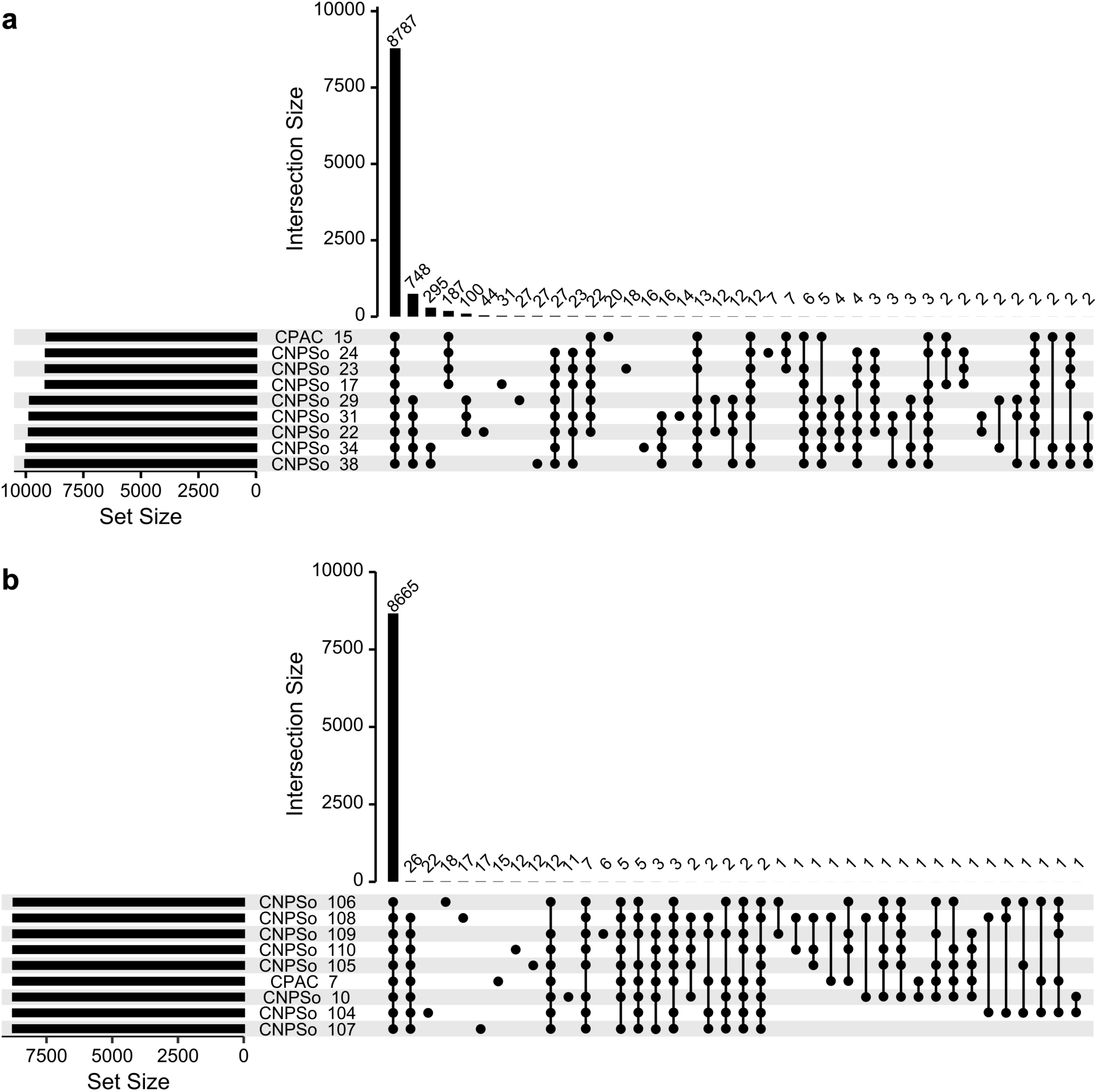
UpSet plot summarizing the pangenome of the reference, parental, and variant strains of the (**a**) *B. japonicum* group or (**b**) *B. diazoefficiens* group. The set size shows the total number of gene families in a given genome, while the intersect size shows the number of gene families conserved across the indicated proteomes.

The *B. diazoefficiens* group has a smaller pangenome than the *B. japonicum* group, consistent with the *B. diazoefficiens* strains being more closely related than the *B. japonicum* strains **(Table 2)**. This pangenome contains a total of 8,665 genes, composed of 8,428 core, 107 accessory, and 130 unique genes. The strain CNPSo 104 presented the largest number of unique genes (22), followed by CNPSo 106 (18), and then CNPSo 107 and CNPSo 108 (each with 17) **(Figure 3b)**. Interestingly, 21% of the dispensable genome fraction falls within SI A, and it includes 37 accessory genes and 13 unique genes **(Table S3)**.

### Nucleotide polymorphisms

We next used Snippy to search for nucleotide variations, using CPAC 15 and CPAC 7 as the reference genomes for the *B. japonicum* and *B. diazoefficiens* groups, respectively. Snippy detects five types of nucleotide variants: single nucleotide polymorphisms (SNPs), multiple nucleotide polymorphisms (MNPs), insertions (ins), deletions (del), and complex variations defined by a combination of SNPs and MNPs. We were particularly interested in nucleotide polymorphisms potentially associated with variation in competitiveness or BNF capacity of the strains **(Table 1)**.

When comparing the sequencing data of the *B. japonicum* strains to the reference strain CPAC 15, a total of 1,150 unique variations were detected. These include 828 SNPs, 21 MNPs, 59 insertions, 89 deletions, and 153 complex variations. Of these, 924 are in protein-coding sequences and 226 in intergenic regions. Excluding synonymous mutations as these are unlikely to have biological effects, 71 variations were detected within genes or intergenic regions of SI A and 66 variations in SI C; no variations were observed in SI B. The variations within the SIs that we predict might be related to competitiveness or BNF are presented in **Table 3**.

**Table 3.**
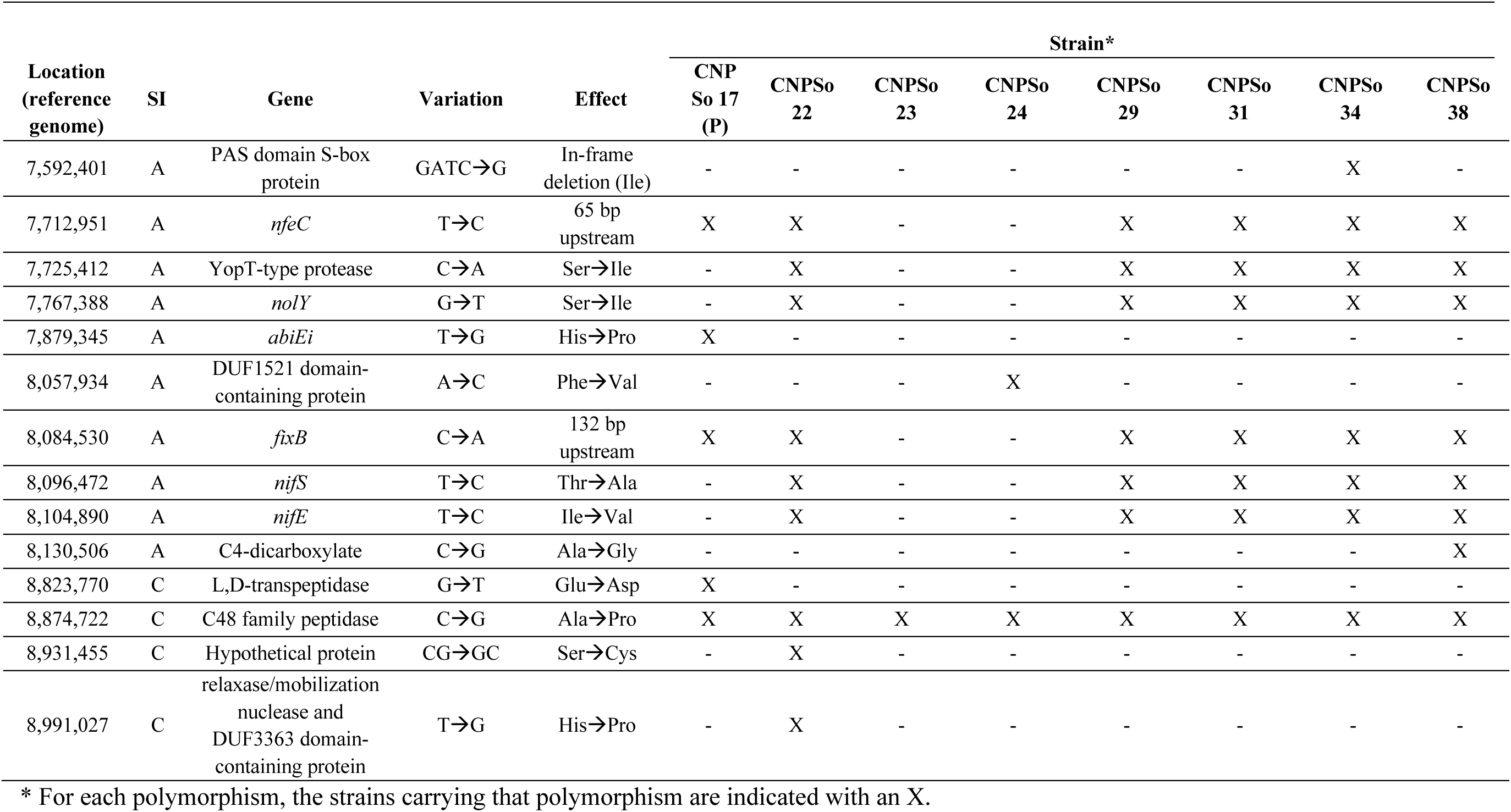
Nucleotide polymorphisms within symbiosis island A potentially related to competitiveness for nodule occupancy or biological nitrogen fixation, which were detected in comparisons between the parental strain (P) or variant strains and the corresponding reference strain, *B. japonicum* CPAC 15.

The strains of the *B. diazoefficiens* group are very closely related, and this conservation is also reflected in the number of SNPs. Within this group, we detected only 57 variations: 24 SNPs, 32 deletions, and one insertion. Of these, 48 are in protein-coding sequences, and nine are in intergenic regions. No MNP or complex variations were identified. Six of the nucleotide variations were detected within SI A, while no variations were observed in SI B or SI C. The variations that we predict might influence the competitiveness or BNF capacity of these strains are shown in **Table 4**.

**Table 4.** Nucleotide polymorphisms within symbiosis island A potentially related to competitiveness for nodule occupancy or biological nitrogen fixation, which were detected in comparisons between the parental strain (P) or variant strains and the corresponding reference strain, *B. diazoefficiens* CPAC 7.

| Location<br>(reference<br>genome) | SI | Gene | Variation | Effect | Strain |  |  |  |  |  |  |  |
| --- | --- | --- | --- | --- | --- | --- | --- | --- | --- | --- | --- | --- |
|  |  |  |  |  | CNPSo 10<br>(P) | CNPSo<br>104 | CNPSo<br>105 | CNPSo<br>106 | CNPSo<br>107 | CNPSo<br>108 | CNPSo<br>109 | CNPSo<br>110 |
| 7,535,266 | A | *nfeC* | GT→G | 108 bp<br>upstream | X | X | X | X | X | X | X | X |
| 7,786,768 | A | pentapeptide<br>repeat domain-containing protein | AC→A | Frameshift<br>deletion | X | X | X | X | X | X | X | X |
| 8,896,771 | A | GNAT family N-acetyltransferase | TG→T | Frameshift<br>deletion | X | X | X | X | X | X | X | X |

\* For each polymorphism, the strains carrying that polymorphism are indicated with an X.

## DISCUSSION

Commercial cropping of soybean began in the southern region of Brazil in the 1950s-1960s, and spread to the Cerrado biome in the central-western region in the 1970s (Hungria and Mendes, 2015). Nowadays, Brazil is the world leader in soybean production, with this production depending on BNF rather than N-fertilizer. The success of BNF in Brazil is due to a long-term program in isolating and selecting highly effective and competitive *Bradyrhizobium* strains adapted to the Brazilian edaphoclimatic conditions and soybean genotypes. *B. japonicum* CPAC 15 and *B. diazoefficiens* CPAC 7 represent 90% of the inoculants currently used in Brazil and have been used since 1992 (Hungria and Mendes, 2015). CPAC 15 is a natural variant of SEMIA 566 (=CNPSo 17) that was isolated from an area of Brazil that had been cropped with soybean and inoculated with CNPSo 17 for more than a decade (Vargas et al., 1992; Hungria and Mendes, 2015). CPAC 7 is a natural variant of SEMIA 586 (=CNPSo 10, =CB 1809), which was isolated following a greenhouse study followed by 2-3 years of adaptation to Cerrado soils (Vargas et al., 1992; Hungria and Mendes, 2015).

In addition to CPAC 15 and CPAC 7, other CNPSo 17 and CNPSo 10 variants were obtained using the same approaches (Boddey and Hungria, 1997). Previous studies have confirmed the parenthood of the variants via rep-PCR profiles (Hungria et al., 1998; Barcellos et al., 2007; Santos et al., 1999), and have identified differences in the phenotypic and symbiotic properties of the variant strains (Boddey and Hungria, 1997; Hungria et al., 1998; Santos et al., 1999; Barcellos et al., 2007). Within the *B. japonicum* group, strains CNPSo 22, CNPSo 23, CNPSo 24, CNPSo 29, CNPSo 31, and CNPSo 34 are more competitive than CPAC 15, while CNPSo 17 and CNPSo 38 show similar competitiveness **(Table 1)** (Hungria et al., 1998). Concerning the *B. diazoefficiens* group, the variants CNPSo 104 and CNPSo 108 show higher BNF capacity than CPAC 7, while CNPSo 105 shows lower BNF capacity, and CNPSo 10 and the other variants display similar BNF capacity to CPAC 7 **(Table 1)** (Santos et al., 1999). Previous studies attempted to identify genes possibly associated with the differences in symbiotic abilities based solely on the genomes or proteomes of CPAC 15, CPAC 7, and the corresponding species type strains (Siqueira et al., 2014; Batista et al., 2010), and more recently, differences in the symbiotic island were examined using draft genome assemblies of the parental strains and a subset of the variants (Bender et al., 2022). We build upon those studies by describing and comparing the complete or high-quality draft genomes of the parental, reference, and variant strains of both groups.

### Symbiotic Island organization

The Cerrado soils are likely a challenging environment due to the high temperatures, long dry-season periods, low pH and nutrient availability, and high aluminum content (Hungria and Mendes, 2015). Consequently, these soils may select for broad genetic and metabolic diversity. HGT is essential for the dissemination of selective advantages and evolution of symbiotic BNF ability among rhizobia, and it contributes to variation in rhizobium-legume symbioses. The symbiotic genes of *Bradyrhizobium* species are usually clustered in symbiotic islands (SIs) located on the chromosome (Ormeño-Orrillo and Martínez-Romero, 2019). Previous work with type strains suggested that the SIs of *B. diazoefficiens* and *B. japonicum* are split into three segments (SIs A, B, and C) with different sizes and locations, and it was suggested this is the result of a larger ancient SI being integrated into the chromosome and then rearranged into separate segments (Kaneko et al., 2002; Kaneko et al., 2011). These same three regions were also identified in the chromosome of *B. japonicum* CPAC 15 and *B. diazoefficiens* CPAC 7 (Siqueira et al., 2014), as well as in all variant and parental strains included in our study.

Consistent with past studies, we observed that SI A contains most of the classical genes required for symbiotic nitrogen fixation. SI A is integrated into a tRNA-valine gene, which is the most common insertion site in bradyrhizobia (Weisberg et al., 2022). SI A carries the *nif* genes encoding proteins responsible for synthesizing and regulating nitrogenase, as well as the *fix* genes involved in oxygen metabolism. SI A also carries the classical genes required for the nodulation process, which include the *nod, noe,* and *nol* genes. In addition, this symbiotic island carries *rhc* and *nop* clusters that encode a T3SS and T3SS effectors (T3Es), respectively, that are involved in other steps of nodulation or alternative nodulation processes (Teulet et al., 2022). The presence of T3SS and T3E genes on the SI of bradyrhizobia has been frequently described (Okazaki et al., 2015; Weisberg et al., 2022), with Teulet et al. (2020) noting that 90% of *Bradyrhizobium* genomes with *nod* genes also encode a T3SS, inferring a common evolutionary origin for both gene groups in this genus. In both the *B. japonicum* and *B. diazoefficiens* groups, the *rhc* and *nop* clusters of all strains correspond to α-RhcI, commonly found in *Nitrobacteraceae* (Teulet et al., 2020).

### B. japonicum plasmids

Although plasmids are unusual in the genus *Bradyrhizobium*, there are some reports of their occurrence. Cyntrin et al. (2008) sequenced the plasmid of the photosynthetic *Bradyrhizobium* sp. BTAi1, a strain able to nodulate aquatic legumes of the genus *Aeschynomene* with no requirement of *nod* genes. Also, up to three plasmids were detected in five *B. japonicum* strains and six *B. elkanii* strains from China, Thailand, and the United States of America (Cyntrin et al., 2008). Whole-genome sequencing carried out by Iida et al. (2015) showed that *B. diazoefficiens* NK6, isolated from root nodules of soybean grown in paddy fields in Niigata (Japan), contains four plasmids (pNK6a, pNK6b, pNK6c and pNK6d), and five other *Bradyrhizobium* strains had plasmids detected by pulsed-field gel electrophoresis. Ormeño-Orrillo and Martínez-Romero (2019), in a study including hundreds of *Bradyrhizobium* genomes, revealed that 35 contained at least one copy of the *repB* gene, indicative of a plasmid. Lastly, *Bradyrhizobium* sp. DOA9 is unique in that its symbiotic genes are found on a symbiotic plasmid rather than on a chromosomal SI; the DOA9 symbiotic plasmid includes nodulation, nitrogen fixation, T3SS, and type IV (T4SS) secretion system genes (Okazaki et al., 2015).

We detected a monophyletic group of five *B. japonicum* variants that each carried two plasmids. Both plasmids were annotated as encoding a number of toxin-antitoxin (TA) systems. These include a putative MazE/MazF TA system, a VapC toxin, and a Phd/YefM antitoxin on plasmid “a”. Similarly, plasmid “b” putatively encodes two VapC family toxins, one VapB family antitoxin, one Phd/YefM family antitoxin, and two nucleotidyl transferase AbiEii/AbiGii toxin family proteins. In addition to functioning as plasmid addition systems, TA systems may also contribute to stress responses, having been linked to cell dormancy, drug-tolerant persister cells, survival during infection, adaptation to hostile environments, and biofilm formation (Schuster and Bertram, 2013; Chen et al., 2021). Interestingly, in *Sinorhizobium meliloti*, a VapB/VapC TA system was implicated in cell growth during symbiotic infection (Arcus et al., 2011). It is tempting to speculate the TA systems of plasmid “a” may contribute to stress tolerance or impact legume symbiosis.

Plasmid “a” carries a T3SS gene cluster that shows only 59% similarity with the T3SS genes of SI A. Other studies also reported the presence of multiple T3SS gene clusters in rhizobia, suggesting that they might be related to host specificity and competitiveness, although further research is required (Teulet et al., 2020, 2022). Although plasmid “b” did not carry genes for a T3SS apparatus, we did detect genes encoding possible T3SS effectors including a C48 family peptidase, an E3 ubiquitin-protein ligase, and a hypothetical protein with an E3 ubiquitin transferase SlrP conserved domain.

Plasmid “a” was also annotated as encoding a HlyD family efflux protein. HlyD belongs to the resistance, nodulation, and cell division (RND) family of efflux transporters. Efflux transporters function as pumps to expel antimicrobial compounds, heavy metals, lipooligosaccharides, proteins, small molecules, and divalent metal cations, and help bacteria to survive in hostile environments. In *Escherichia coli*, HlyD is responsible for secreting hemolysin in pathogenic infections (Zgurskaya and Nikaido, 2000). Likewise, we detected genes putatively encoding efflux transporter proteins on plasmid “b”, including two HlyD family secretion periplasmic adaptor subunit, and a HlyB protein family. It is possible that these efflux proteins improve the competitiveness of bradyrhizobia, by increasing resistance to antimicrobial compounds produced by other soil microbes or during the symbiotic association with legumes (Maximiano et al., 2021).

Several other proteins related to stress responses were identified on the plasmids, all of which may help bradyrhizobia better survive the challenging conditions faced in Cerrado soils. Plasmid “a” carries a gene putatively encoding a molecular chaperone (HtpG), while plasmid “b” carries genes putatively encoding a cold-shock protein of the CspA family, a GroEL chaperonin, and the co-chaperone GroES. The chaperone HtpG is a heat shock protein involved in maintaining protein-folding homeostasis in *E. coli* under high-temperature conditions (Thomas and Baneyx, 2000), and was up-regulated during salt stress in *Rhizobium tropici* (Maximiano et al., 2021). The CspA family includes RNA chaperones responsible for regulating the expression of target genes during temperature downshifts (Alexandre and Oliveira, 2013). GroEL and its cofactor GroES represent a chaperone system constitutively expressed under normal conditions and is important for bacterial protein folding by creating a hydrophilic environment. These proteins are upregulated in heat stress conditions, preventing protein denaturation (Alexandre and Oliveira, 2013). *Bradyrhizobium* strains often carry *groEL-groES* operons; Batista et al. (2010) detected two spots of GroEL proteins in the *B. japonicum* CPAC 15 proteome, while Gomes et al. (2014) found GroEL spots in the *B. diazoefficiens* CPAC 7 proteome.

In addition to the chaperones, plasmid “a” putatively encodes a trehalose-6-phosphate synthase (OtsA), while plasmid “b” putatively encodes a choline dehydrogenase (BetA). Trehalose-6-phosphate synthases are involved in the biosynthesis of trehalose, a nonreducing disaccharide involved in bacterial tolerance against desiccation, heat, cold, oxidation, and osmotic stresses (Ledermann et al., 2021). OtsA orthologs have been linked to enhanced nodule occupancy competitiveness of *B. diazoefficiens* via improved osmotic tolerance in the early stages of soybean nodulation (Sugawara et al., 2009; Ledermann et al., 2021). Similarly, OtsA was linked to both free-living osmoadaption in *S. meliloti* and competitiveness for alfalfa nodulation (Domínguez-Ferreras et al., 2009). Likewise, BetA is involved in osmoadaptation through production of the osmoprotectant glycine betaine, and its disruption in *S. meliloti* prevents this organism from using environmental choline as an osmoprotectant (Pocard et al., 1997). The *S. meliloti bet* operon is also highly induced in alfalfa nodules (Mandon et al., 2003). Considering the edaphoclimatic conditions of the Cerrado region, which has year-round high temperatures and long dry-season periods that could result in osmotic stresses, the plasmid-encoded chaperons, OtsA, and BetA may improve the fitness of bradyrhizobia in Cerrado soils.

We also detected a putative O-antigen biosynthesis gene cluster on plasmid “b”, which could impact the O-antigen structure of the variants carrying this plasmid. The O-antigen comprises repeating oligosaccharides units, and together with lipid A and the core oligosaccharide, composes the lipopolysaccharides (LPS) of Gram-negative bacterial cell walls. O-antigens are structurally diverse and are essential for bacteria-host interactions, by suppressing host defenses. We identified a full pathway for the production of dTDP-L-rhamnose, a common component of bacterial O-antigens. In addition, we identified two glycosyltransferase family-2 proteins, two glycosyltransferase family-4 proteins, an ABC transporter permease, an ABC transporter ATP-binding protein, two glycosyltransferases, and an acetyltransferase that may also be related to O-antigen biosynthesis (Reeves et al., 1996).

Overall, we hypothesize that plasmids “a” and “b” may provide adaptative advantages to the strains carrying these mobile elements under the hostile conditions faced in Cerrado soils. By improving their saprophytic ability in the soil, the strains would also be more competitive for nodule occupancy.

### Variation in the gene content of the symbiotic islands

Whole genome alignments indicated high conservation of gene order across the chromosomes within the *B. japonicum* and *B. diazoefficiens* genomes. Although we detected the same possible inversion in *B. japonicum* CPAC 15 and CNPSo 22 (which are not sister taxa), this may instead reflect assembly errors as these were the only two genomes not assembled using Flye. Similarly, the SIs A, B, and C of *B. japonicum* and *B. diazoefficiens* were also highly syntenic within each group, despite these regions being enriched for IS and transposase genes (Barros-Carvalho et al., 2018).

To further evaluate how genomic variation may be correlated with the phenotypic differences of the variants, pangenome analyses were performed to identify variation in gene content across strains. We focused on variations in SI A, as this region contains the primary genes required for symbiotic nitrogen fixation. The *B. japonicum* strains carrying plasmids have a SI A smaller than those that lack plasmids. This difference is driven by a contiguous region of ∼50 genes present in the plasmidless strains, which is replaced by a set of 27 genes in the plasmid-containing strains. In the strains lacking plasmids, about half of the 50 genes are annotated as hypothetical genes, while the others included genes encoding type II and type IV secretion systems proteins, oxidoreductases, transcriptional regulators, IS, and transposases **(Table S2)**. In the plasmid-containing strains, the 27 genes include eight hypothetical genes, transposases, and genes encoding proteins that may influence their saprophytic ability, such as a cold shock protein, carbohydrate and peptide transport systems, and a LuxR-like transcriptional regulators **(Table S2)**.

A few other interesting differences were observed within the *B. japonicum* group **(Table S2)**. The parental strain CNPSo 17 appears to have lost a NoeE-like protein, which is a sulfotransferase related to modifications in the Nod factors and host specificity (Wang et al., 2019). Interestingly, CNPSo 38, which is less competitive than most variants of the *B. japonicum* group, gained a gene putatively encoding a second copy NopM, an effector protein secreted by the T3SS and that is related to negative effects in the interaction with legumes (Kambara et al., 2009). On the other hand, no gain or loss of a specific gene that might explain the higher competitiveness of CNPSo 22, CNPSo 23, CNPSo 24, CNPSo 29, CNPSo 31, and CNPSo 34 from the *B. japonicum* group was identified.

Likewise, several interesting differences were observed within the *B. diazoefficiens* group within SI A **(Table S3)**. CNPSo 106, a variant with equal BNF capacity and competitiveness to CPAC 7, lost a gene coding for acetyltransferase containing a GNAT domain. Acetyltransferases are involved in Nod factors biosynthesis (Wang et al., 2019) and, recently, a GNAT acetyltransferase was identified as related to competitiveness for *Pisum sativum* in *R. leguminosarum* bv. *viciae* (Boivin et al., 2019). In addition, CNPSo 106 may have lost putative *nopM* and a *bacA*-like genes. The negative impact of *nopM* was described above, while *bacA* and *bclA* (*bacA*-like) encode peptide transporters essential for symbiosis with legumes that produce nodule-specific cysteine-rich (NCR) peptide, but not for symbiosis with legumes such as soybean that do not produce NCR peptides (Glazebrook et al., 1993; Karunakaran et al., 2010; Guefrachi et al., 2015). Interestingly, CNPSo 104, the most competitive and efficient nitrogen-fixer of this group, appears to have lost nine genes encoding five transposases, three hypothetical proteins, a sulfite exporter TauE/SafE family protein, and a sulfurtransferase important to sulfur and carbon cycles, while gaining two hypothetical genes and two putative transposases. We did not observe any obvious gene gains or losses in SI A related to the higher BNF capacity of CNPSo 104 and CNPSo 108 compared to the rest of the *B. diazoefficiens* group.

### Nucleotide variations within SI A of the *B. japonicum* group

In addition to examining variation in gene presence/absence, we compared the genomes of the variant and parental strains to identify nucleotide sequence variations potentially associated with differences in BNF efficiency or competitiveness for nodule occupancy. Recently, Bender et al. (2022) analyzed SNPs in the SI A of some of the same strains used in our study; while some SNPs were detected in both studies, others were not, likely as we included more strains, used complete genomes, and used different annotation and analysis tools.

Several interesting nucleotide polymorphisms were detected within SI A of the *B. japonicum* group of strains. We detected a SNP in a gene encoding an AbiEi family antitoxin found only in the parental strain CNPSo 17, which has equal competitiveness but lower BNF capacity than CPAC 15 (**Table 1**). A study by Chen et al. (2021) demonstrated that mutation of the *abiEi* antitoxin gene of *Mesorhizobium huakuii* did not alter the number of nodules but strongly affected bacteroid occupancy and BNF efficiency. In addition, 21 genes directly related to symbiotic BNF were up- or down-regulated in the transcriptome of the *M. huakuii abiEi* mutant. We therefore hypothesize that the nucleotide variation detected in *abiEi* might negatively affect the BNF capacity of the parental strain.

Strain CNPSo 22 contained a unique SNP in a hypothetical gene containing a conserved domain of the extra-cytoplasmic function (ECF) σ factor of the *rpoE* gene. The σ factors are ubiquitous in bacterial genomes and are involved in the control of gene expression by binding to RNA polymerase. A putative ECF σ factor of *S. meliloti* is associated with several stress conditions including heat and salt stress, as well as carbon and nitrogen starvation (Sauviac et al., 2007). In addition, Martínez-Salazar et al. (2009) suggested that *rpoE4* of *Rhizobium etli* is a general regulator involved with saline and osmotic responses, oxidative stress, and cell envelope biogenesis. Gourion et al. (2009) showed that *B. diazoefficiens* USDA 110^T^ ECF σ factor mutants are more sensitive to heat and desiccation upon carbon starvation than the wild type. In addition, mutants formed nodules with reduced number, size, and BNF capacity in association with *G. max* and *Vigna radiata*, suggesting that ECF σ factors are important for *Bradyrhizobium* symbiosis (Gourion et al., 2009). Considering that CNPSo 22 is a highly competitive variant, we hypothesize that the nucleotide variation in the ECF σ factor gene of SI A may be a contributing factor.

Strain CNPSo 24 has higher competitiveness and efficiency of BNF than CPAC 15, and it contains a unique SNP on in a gene encoding a DUF1521 domain-containing a protein homolog to the T3E NopE1. Zenher et al. (2008) detected NopE1 in mature *Macroptilium atropurpureum* nodules hosting *B. japonicum* associated, indicating a putative function of this effector in rhizobia-legume symbiosis. Wenzel et al. (2010) also identified NopE1 in nodules and showed that mutation of *B. japonicum nopE1* and its homolog, *nopE2*, results in a reduced number of nodules on *M. atropurpureum* and *G. max*. However, the same double mutant significantly increased the number of nodules in *V. radiata*, suggesting the impact is host specific (Wenzel et al., 2010). The DUF1521 domain-containing protein of CNPSo 24 is located three genes downstream to several other *rhc* genes of SI A and thus may play a role in symbiotic BNF; however, whether this variation positively or negatively influences the symbiotic capacity of CNPSo 24 remains to be evaluated.

Strain CNPSo 34 presents higher competitiveness but lower BNF capacity than CPAC 15. CNPSo 34 contains an in-frame deletion in a gene encoding a PAS domain S-box protein. Prokaryotic PAS domains usually are part of two-component regulatory systems composed of a histidine kinase sensor and a response regulator. Several BNF and nodulation proteins have a PAS domain that serves as an oxygen and/or redox sensor, which are important for nitrogenase activity and energy metabolism, respectively (Taylor and Zhulin, 1999). Examples include FixL of the FixL/FixJ two-component system that detects environmental oxygen levels and regulates expression of BNF genes (Gilles-Gonzales and Gonzales, 1993). Besides FixL/FixJ, *Azorhizobium caulinodans* also has also the NtrY/NtrX two-component system; NtrY is a membrane-associated sensor with a PAS domain, which may be involved in sensing extracellular nitrogen levels (Taylor and Zhulin, 1999). NifU has a PAS domain at the N-terminus, possibly related to iron and sulfur mobilization for the iron-sulfur cluster of nitrogenase (Taylor and Zhulin, 1999). In addition, the *nodV/nodW* genes are involved in regulating the nodulation genes through flavonoid signals; NodV has four PAS domains (Taylor and Zhulin, 1999). Given that the focal PAS domain S-box protein is found within SI A, we hypothesize that it is also related to symbiosis, and that the in-frame deletion in CNPSo 34 contributes to its symbiotic phenotypes.

Strain CNPSo 38, with equal competitiveness and lower BNF capacity than CPAC 15, carried a unique SNP in *dctA*. The *dctA* gene is essential for symbiotic nitrogen fixation as it encodes a transporter responsible for transporting the C_4_-dicarboxylates malate, succinate, and fumarate, which are the primary carbon sources received by rhizobia in nodules (Ronson et al., 1984; Finan et al., 1983). Therefore, as *dctA* is essential for symbiotic nitrogen-fixation, the nucleotide variation within this gene may negatively impact the BNF capacity of CNPSo 38, and potentially also its saprophytic ability.

The monophyletic group of plasmid-carrying strains CNPSo 22, CNPSo 29, CNPSo 31, CNPSo 34, and CNPSo 38 carry a SNP in a gene encoding a YopT-type cysteine protease, an effector protein usually found in the pathogenic bacteria *Pseudomonas syringae* and *Yersina* (Kambara et al., 2009), and homologous to the T3E NopT. *S. fredii* NGR234 *nopT* mutants show improved nodulation with *Phaseolus vulgaris* and *Tephrosia vogelii*, and are negatively impacted in their association with *Crotalaria juncea* (Kambara et al., 2009). Conversely, *Bradyrhizobium vignae* ORS3257 *nopT* mutants form fewer nodules on *V. unguiculata* and *V. mungo* (Songwattana et al., 2021). We therefore hypothesize that the SNP in the gene encoding a YopT-type cysteine protease may impact nodulation and deserves further studies. Interestingly, this group of strains also contain a SNP located in *nolY* encoding an isoflavone *nodD*-dependent protein related to infection events. *B. diazoefficiens* USDA 110^T^ *nolY* mutants show a slight nodulation defect on *G. max*, *M. atropurpureum,* and *V. unguiculata*, and a severe nodulation defect on *V. mungo* (Dockendorff et al., 1994). These five variant strains also carry SNPs in two genes related to nitrogenase biosynthesis, *nifS* and *nifE*. NifS is a cysteine desulfurase involved in donating sulfur for the FeS metallocluster of Fe-protein of nitrogenase (Ludden, 1993). NifE participates in the Fe-Mo-co metallocluster synthesis of the MoFe-protein of nitrogenase, along with the *nifB, nifH, nifN, nifQ,* and *nifV* genes (Kennedy and Dean, 1992). These two SNPs were also detected in CNPSo 22 and CNPSo 38 by Bender et al. (2022) and could impact the BNF efficiency of these variant strains.

All variants with plasmids and the parental strain CNPSo 17 carry a SNP in an intergenic region 65 bp upstream of a gene encoding the nodule efficiency protein C (*nfeC*) of SI A. The nodule efficiency proteins were first identified in *S. meliloti* GR4 and are associated with improved nodulation efficiency and competitiveness with *Medicago* (Sanjuan and Olivares, 1989). Similarly, deletion of *nfeC* in *B. diazoefficiens* USDA 110^T^ resulted in delayed nodulation on soybean and reduced competitiveness for nodule occupancy (Chun and Stacey, 1993). This group of strains also have a SNP in an intergenic region 132 bp upstream of a gene encoding an electron transfer flavoprotein alpha subunit FixB family protein in SI A. In addition to *nod* and *nif* genes, the *fix* genes are important to BNF as they encode electron transfer proteins that function under microaerobic conditions (Earl et al., 1987). The nucleotide variations upstream of *nfeC* and *fixB* may alter their expression and consequently impact nodulation and nitrogen fixation, respectively. The reference strain CPAC 15 has a unique SNP in a gene putatively encoding a C48-family peptidase. Young et al. (2010) suggested that the Bll8244 protein of *B. diazoefficiens* USDA 110^T^, a homolog of C48-family peptidases, functions as a genistein secreted T3E protein. The C48-family protein of our focal strains contains a conserved domain of small ubiquitin-like modifier (SUMO) proteases, the main effector family found in the genus *Bradyrhizobium* (Teulet et al., 2020). Moreover, a SUMO domain was identified in a putative effector protein of *B. japonicum* Is-34 that is responsible for the inability of this strain to nodulate the soybean *Rj4* genotype (Tsurumaru et al., 2010). Consequently, we hypothesize that the SNP in the focal gene encoding a C48-family peptidase in CPAC 15 may impact competitiveness and nodulation, and, therefore, it should be carefully investigated in further studies.

### Nucleotide variations within SI A of the *B. diazoefficiens* group

The reference strain CPAC 7 carried three interesting nucleotide polymorphisms in SI A compared to the rest of the *B. diazoefficiens* strains. CPAC 7 contained a one nucleotide insertion 108 bp upstream to *nfeC*. As discussed above, *nfeC* is related to nodulation and competitiveness of rhizobia. We hypothesize that the nucleotide insertion may impact expression, and thus competitiveness, of CPAC 7. CPAC 7 also contains a one nucleotide frameshift insertion in a gene putatively encoding a pentapeptide repeat-containing protein homologous to YjbI, a truncated hemoglobin of *Bacillus subtilis*. Rogstam et al. (2007) demonstrated that a *B. subtilis yjbI* mutant is hypersensitive to sodium nitroprusside, a source of nitric oxide. Therefore, the CPAC 7 variation within *yjbI* of the *B. diazoefficiens* group may impact the saprophytic ability of the strains. Lastly, CPAC 7 contains a one nucleotide frameshift insertion in a gene putatively encoding a N-acetyltransferase GNAT family. As N-acetyltransferases may promote modifications of Nod factors (Wang et al., 2019), this mutation could impact competitiveness for nodulation and host specificity.

### Conclusions

Using whole genome sequencing and comparative genomic methodologies, we identified numerous genomic differences across the *B. japonicum* and *B. diazoefficiens* variants. Overall, a higher amount of variability was found within the *B. japonicum* strains compared to the *B. diazoefficiens* group. Interestingly, we detected a remarkable diversity of mobile elements in the *B. japonicum* group, with a high number of insertion sequences that may have contributed to genome rearrangements and HGT. The *B. japonicum* group has a comparatively large pangenome, and a total of 1,150 nucleotide polymorphisms were detected across the strains, including 71 non-synonymous variations within SI A. In addition, a monophyletic group of five *B. japonicum* variants unexpectedly carry two plasmids, with several of the plasmid-encoded genes putatively associated with tolerance to environmental stresses. Less variation was observed in the *B. diazoefficiens* group. The chromosomes and SIs were highly syntenic, and a smaller pangenome was detected in comparison to the *B. japonicum* group. In total, only 57 nucleotide polymorphisms were detected, of which only six are located in SI A. No plasmids were detected in the *B. diazoefficiens* strains. The large difference in level of genetic variability of the two groups likely results from how the variants were originally isolated. The *B. japonicum* variants were isolated as highly competitive variants following more than ten years of growth of CPAC 15 in Cerrado soils. In contrast, the *B. diazoefficiens* variants were selected as strains with high BNF capacity and were selected based colony morphological differences followed by only a few years of adaptation to Cerrado soils (Hungria and Mendes, 2015).

In conclusion, we identified numerous genetic variations – including gene gains/losses, plasmid acquisition, and nucleotide polymorphisms – across natural variants of the soybean inoculants *B. japonicum* CNPSo 17 and *B. diazoefficiens* CNPSo 10, highlighting the high plasticity of *Bradyrhizobium* genomes. The level of genetic variability correlated with the length of time the parental strains were allowed to adapt to their new environment. We hypothesize that many of the genetic variations reflect early adaptation to the stressful conditions of Cerrado soils that might improve saprophytic ability, or that alter competitiveness or BNF capacity with local soybean genotypes. In general, single genomic differences able to explain the phenotypic differences of the variants were not obvious, suggesting that the observed alterations in competitiveness for nodule occupancy and BNF capacity may instead reflect the cumulative impact of multiple genomic variations.

## MATERIALS AND METHODS

### Bradyrhizobium strains

This study examined 18 *Bradyrhizobium* strains, nine belonging to the species *B. japonicum* and nine to the species *B. diazoefficiens*. For each species, the strains included one parental genotype previously used as a commercial inoculant in Brazil, one natural variant (the reference genome) used in commercial inoculants from 1992 until the present, and seven other natural variants. The strains used in this study, as well as their BNF and competitiveness capacities, are shown in **Table 1**. The parenthood of the strains within each group was confirmed by their BOX-PCR profiles (**Figure S3**). The background of the strains is detailed in the discussion. All strains are deposited in the ‘Diazotrophic and Plant Growth Promoting Bacteria Culture Collection of Embrapa Soybean’ (WFCC Collection # 1213, WDCM Collection # 1054), Londrina, Paraná State, Brazil.

### DNA extraction and genome sequencing

A high-quality draft genome of CPAC 15 was previously sequenced by Siqueira et al. (2014) and was retrieved from the National Center of Biotechnology Information (NCBI; GenBank accession CP007569). Draft genomes of strains CNPSo 17, CNPSo 22, CNPSo 23, CNPSo 38, CNPSo 10, CNPSo 104, CNPSo 105, and CNPSo 107 were previously reported by Bender et al., (2022) and their raw Illumina data was used in the process of completing their genomes in this study. All other strains were sequenced as part of this study. Strains were grown on a modified-yeast mannitol (YM) medium (Hungria et al., 2016) at 28℃ for five days. The total DNA of each strain was extracted using the DNeasy Blood and Tissue kit (Qiagen), according to the manufacturer’s instructions.

Libraries for Illumina sequencing were prepared following the instructions of the Nextera XT kit (# 15031942 v01) and sequenced on a MiSeq instrument to generate 300 bp paired-end reads. ONT sequencing was performed using a Rapid Barcoding Kit (SQK-RBK004) and an R9.4.1 flow cell on a minION device. ONT basecalling and demultiplexing were performed using Guppy version 5.011+2b6dbffa5 and the high accuracy model (ONT). Sequencing statistics are provided in **Table 2**.

### Genome assembly

The quality of the Illumina reads was checked using the FASTQC tool (bioinformatics.babraham.ac.uk/projects/fastqc/). Subsequently, adapter sequences and low-quality bases were trimmed using trimmomatic version 0.39 (Bolger et al., 2014) with the parameters: LEADING:3 TRAILING:3 SLIDINGWINDOW:4:15 MINLEN:36 ILLUMINACLIP:NexteraPE.fa:2:30:10. The phiX sequences were removed using the run_bowtie2_subtract_unmapped_reads.pl script (github.com/tomdeman-bio/Sequence-scripts) with the dependencies bowtie2 version 2.4.5 (Langmead et al., 2012) and samtools version 1.15.1-33-g906f657 (Danecek et al., 2021).

De novo genome assembly was performed using Flye version 2.9-b1768 (Kolmogorov et al. 2019) with the ONT reads. Assemblies were checked for overlaps between the two ends of a contig using NUCmer version 4.0.0rc1 (Kurtz et al., 2004), and if identified, one copy of each overlap was removed. The assemblies were first polished with the ONT reads using Racon version 1.4.13 (Vaser et al., 2017) followed by Medaka version 1.4.1 (github.com/nanoporetech/medaka); read mapping was performed with Minimap2 version 2.20-r1061 (Li, 2018). The assemblies were then polished with the Illumina reads using Pilon version 1.24 (Walker et al., 2014) and Racon; read mapping was performed using bwa version 0.7.17-r1198-dirty (Li and Durbin, 2009). NUCmer was used again to check for overlaps between two ends of a contig, with one copy removed if found.

As Flye did not produce a fully circular chromosome for strains CNPSo 22 and CNPSo 38, *de novo* assembly was repeated using Unicycler version 0.5.0 (Wick et al., 2017) with both the ONT and Illumina reads. The Flye and Unicycler assemblies were then merged using the patch function of RagTag version 2.1.0 (Alonge et al., 2019). The procedure resulted in a circular chromosome for strain CNPSo 22, and the resulting assembly was used for downstream analyses. As this process did not improve the quality of the CNPSo 38 genome, we continued with the Flye-based assembly for subsequent steps.

Lastly, the assemblies were reoriented such that the chromosomes began at the putative origin of replication (Kaneko et al., 2011), using circlator version 1.5.5 with the fixstart option (Hunt et al., 2015). The genome of *B. japonicum* CPAC 15 was similarly reoriented.

### Genome annotation

Genome assemblies were annotated using the NCBI Prokaryotic Genome Annotation Pipeline (PGAP) program version 2022-02-10.build5872 (Tatusova et al. 2016). The symbiotic island (SI) regions A and B were detected as described by Weisberg et al. (2022), whereas SI region C was detected according to Kaneko et al. (2011).

### Genome assembly statistics and average nucleotide identity

The quality of the genome assemblies was evaluated using the web-based tool QUAST: Quality Assessment Tool for Genome Assemblies (Gurevich et al., 2013). Pairwise average nucleotide identity (ANI) was calculated using FastANI version 1.33 with default parameters (Jain et al., 2018). For ANI calculations, the genomes of *B. japonicum* USDA 6^T^ (GenBank accession NC_017249.1) and *B. diazoefficiens* USDA 110^T^ (Genbank accession NC_004463.1) were downloaded from NCBI.

### Pangenome calculation

Prior to pangenome analysis, the genome sequences were reannotated using Prokka version 1.14.6 (Seemann, 2014) to produce GFF3 files compatible with Roary. The GFF3 files produced by Prokka were used as input for Roary version 3.11.2 (Page et al. 2015) using the default identity threshold of 95%. Pangenomes were calculated separately for each group of strains (*B. japonicum* and *B. diazoefficiens*). The pangenomes were visualized as UpSet plots using the R package UpSetR version 1.4.0 (Conway et al., 2017).

### Strain phylogenetic analysis

A pangenome of all 18 strains, as well as of *B. japonicum* USDA 6^T^ and *B. diazoefficiens* USDA 110^T^, was constructed using Roary with the -e and -n options to produce a concatenated alignment of the 2,689 core genes; the alignment was produced using MAFFT version 7.310 (Katoh and Standeley, 2013). The alignment was trimmed with trimAl version 1.4rev22 (Capella-Gutierréz et al., 2009) with the automated1 option, and used to construct a maximum likelihood phylogeny with RAxML version 8.2.12 (Stamatakis, 2014), under the GTRCAT model with 1,000 bootstrap inferences. The phylogeny was visualized using iTOL (Letunic and Bork 2016).

### Nucleotide variant identification

Single nucleotide polymorphisms (SNPs) and insertion/deletions (INDELs) were identified using Snippy version 4.6.0 (github.com/tseemann/snippy) and the PGAP output, using CPAC 7 and CPAC 15 as the reference genomes. Polymorphisms located in intergenic regions were visualized with the Integrative Genomics Viewer (IGV) program version 2.11.9 (Robinson et al., 2011) to identify polymorphisms in the promoter regions of genes related to competitiveness, BNF capacity, and saprophytic ability. The protein sequences of genes upstream or downstream of the variation, as well as the protein sequences of genes located in the SIs containing variations, were annotated using the NCBI conserved domain database (CDD) version 3.17 (Lu et al., 2020) with the CD-search option (Marchler-Bauer and Bryant, 2004) to identify functional units in the protein sequences.

### Genome synteny analyses

The genome sequences of the *B. japonicum* and *B. diazoefficiens* groups were compared using BLASTn from BLAST+ version 2.14.0 (Altschul et al., 1990) with the following parameters: identity threshold of 98%, maximum number of high scoring pairs (HSP) of 200, and a minimum raw gapped score of 10,000. The output files were used as input for the Artemis Comparison Tool (ACT) version 18.2.0 (Carver et al. 2005) to visualize genome synteny. The SI A, B, and C of each group were selected using the program faidx version 0.7.2.1 according to the delimitations established by Weisberg et al. (2022) and Kaneko et al. (2010), and ACT comparisons were performed as described above.

### Data availability

The genome assemblies of this study are available on NCBI BioProject database under the accessions numbers PRJNA1026581 for *B. japonicum* strains and PRJNA1026967 for *B. diazoefficiens* strains. PGAP and prokka annotations and code used for the analysis reported in this study are available on GitHub at https://github.com/MSKlepa/Bradyrhizobium_variants.

## ACKNOWLEDGEMENTS

We gratefully acknowledge Leonardo Araujo Terra for helpful bioinformatics advice, and Sabhjeet Kaur for assistance with the ONT sequencing. The group of M.H. is supported by the INCT project “Plant Growth-Promoting Microorganisms for Agricultural Sustainability, Environmental Responsibility” (CNPq 465133/2014-4, Fundação Araucária-STI 043/2019). Research in the G.C.D. laboratory is supported by a Natural Sciences and Engineering Research Council of Canada (NSERC) Discovery Grant. M.S.K acknowledges the Coordenação de Aperfeiçoamento de Pessoal de Nível Superior (CAPES), Finance Code 001, for the PDSE fellowship. M.S.K was further supported by a Mitacs Globalink Research Award and a scholarship from Fundação Araucária.

## Notes

https://github.com/MSKlepa/Bradyrhizobium_variants

### Competing Interest Statement

The authors have declared no competing interest.

