## Supplementary Material for "Comparative genomic analysis of *Bradyrhizobium* strains with natural variability in the efficiency of nitrogen fixation, competitiveness, and adaptation to stressful edaphoclimatic conditions"

**This PDF file includes:**

Figures S1 to S3 (pages 2-4)
Tables S1 to S3 (pages 5-24)

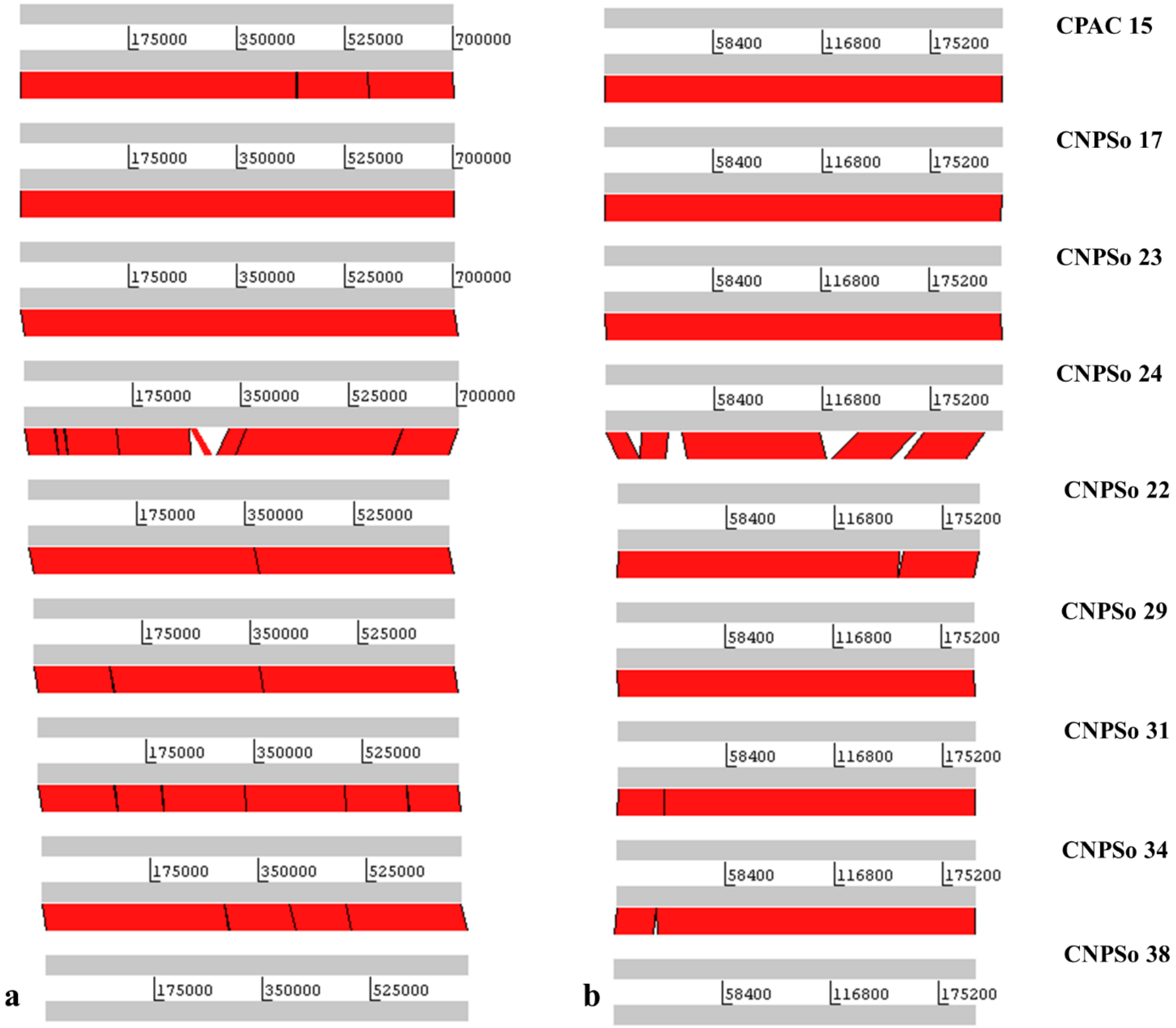


**Figure S1.** Synteny analysis of (**a**) symbiosis island A or (**b**) symbiosis island C of the *B. japonicum* parental strain CNPSo 17, reference strain CPAC 15, and other variants. Pairwise alignments were performed, and homologous regions between pairs of genomes are connected by red lines (if in the same orientation) or blue lines (if in the inverse orientation).

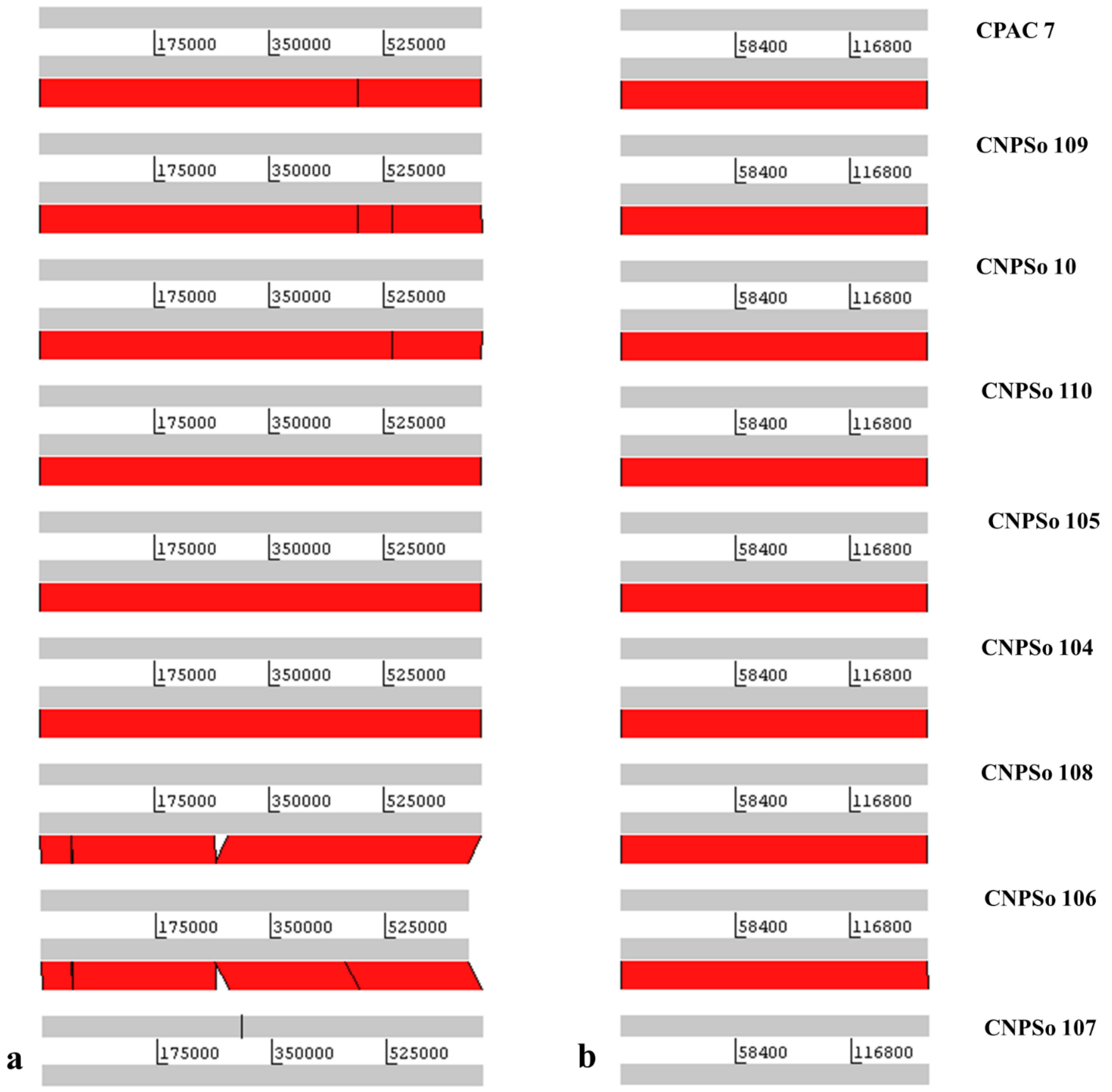


**Figure S2.** Synteny analysis of (**a**) symbiosis island A or (**b**) symbiosis island C of the *B. diazoefficiens* parental strain CNPSo 10, reference strain CPAC 7, and other variants. Pairwise alignments were performed, and homologous regions between pairs of genomes are connected by red lines (if in the same orientation) or blue lines (if in the inverse orientation).

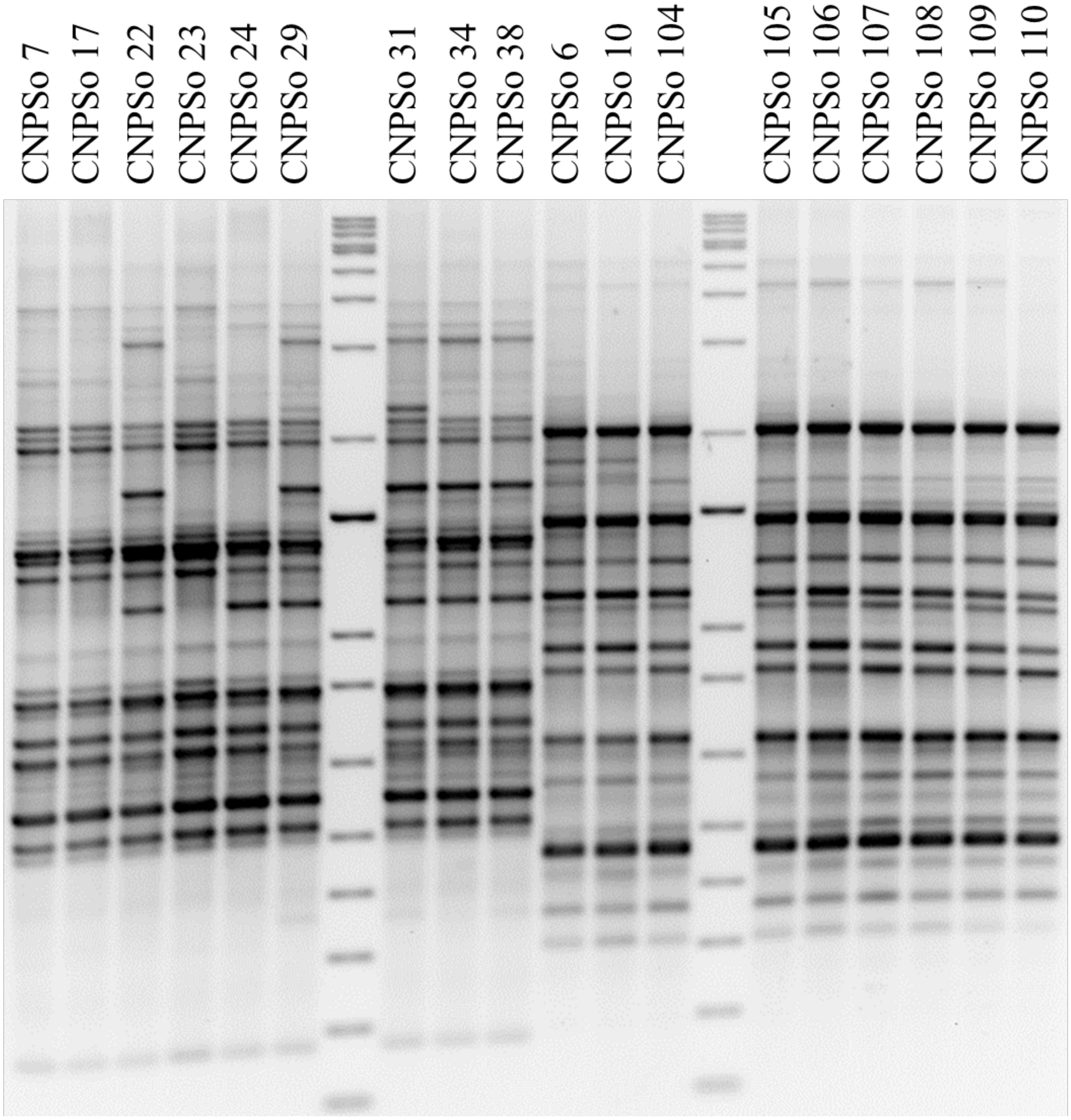


**Figure S3.** BOX-PCR profiles of the parental, reference, and variant strains of the *B. japonicum* and *B. diazoefficiens* groups.

**Table S1.** Gene content of plasmids "a" and "b" of CNPSo 22, CNPSo 29, CNPSo 31, CNPSo 34 and CNPSo 38 of the *B. japonicum* group.

| Annotation | Conserved domain/protein | pBjCNPSo22a | pBjCNPSo29a | pBjCNPSo31a | pBjCNPSo34a | pBjCNPSo38a |
|---|---|---|---|---|---|---|
| type II toxin-antitoxin system Phd/YefM family antitoxin | - | pgaptmp_009680 | pgaptmp_000296 | pgaptmp_009645 | pgaptmp_009804 | pgaptmp_009824 |
| GNAT family N-acetyltransferase | - | pgaptmp_009707 | pgaptmp_000323 | - | - | - |
| antitoxin MazE family protein | - | pgaptmp_009711 | pgaptmp_000327 | pgaptmp_009676 | pgaptmp_009835 | pgaptmp_009855 |
| type II toxin-antitoxin system PemK/MazF family toxin | - | pgaptmp_009712 | pgaptmp_000328 | pgaptmp_009677 | pgaptmp_009836 | pgaptmp_009856 |
| HlyD family efflux transporter periplasmic adaptor subunit | - | pgaptmp_009726 | pgaptmp_000342 | pgaptmp_009691 | pgaptmp_009850 | - |
| glycosyltransferase | CelA, BscA | pgaptmp_009727 | pgaptmp_000343 | pgaptmp_009692 | pgaptmp_009851 | pgaptmp_009870 |
| molecular chaperone HtpG | - | pgaptmp_009729 | pgaptmp_000345 | pgaptmp_009694 | pgaptmp_009853 | pgaptmp_009872 |
| hypothetical protein | Ulp1 protease family | pgaptmp_009732 | pgaptmp_000348 | pgaptmp_009697 | pgaptmp_009856 | pgaptmp_009875 |
| LysR substrate-binding domain-containing protein | nitrogen assimilation transcriptional regulator | pgaptmp_009759 | pgaptmp_000375 | pgaptmp_009724 | pgaptmp_009883 | pgaptmp_009902 |
| EscF/YscF/HrpA family type III secretion system needle major subunit | - | pgaptmp_009760 | pgaptmp_000376 | pgaptmp_009725 | pgaptmp_009884 | pgaptmp_009903 |
| hypothetical protein | type III secretion apparatus protein, YscD/HrpQ family | pgaptmp_009761 | pgaptmp_000377 | pgaptmp_009726 | pgaptmp_009885 | pgaptmp_009904 |

| | | | | | | |
|---|---|---|---|---|---|---|
| EscU/YscU/HrcU family type III secretion system export apparatus switch protein | StcU | pgaptmp_009775 | pgaptmp_000391 | pgaptmp_009740 | pgaptmp_009901 | pgaptmp_009919 |
| type III secretion system export apparatus subunit SctT | - | pgaptmp_009776 | pgaptmp_000392 | pgaptmp_009741 | pgaptmp_009902 | pgaptmp_009920 |
| type III secretion system export apparatus subunit SctS | - | pgaptmp_009777 | pgaptmp_000393 | pgaptmp_009742 | pgaptmp_009903 | pgaptmp_009921 |
| type III secretion system export apparatus subunit SctR | - | pgaptmp_009778 | pgaptmp_000394 | pgaptmp_009743 | pgaptmp_009904 | pgaptmp_009922 |
| FliM/FliN family flagellar motor switch protein | SctQ | pgaptmp_009779 | pgaptmp_000395 | pgaptmp_009744 | pgaptmp_009905 | pgaptmp_009923 |
| hypothetical protein | type III protein | pgaptmp_009785 | pgaptmp_000401 | pgaptmp_009750 | pgaptmp_009911 | pgaptmp_009929 |
| trehalose-6-phosphate synthase | OtsA | pgaptmp_009787 | pgaptmp_000403 | pgaptmp_009752 | pgaptmp_009913 | pgaptmp_009931 |
| type III secretion protein | | pgaptmp_009793 | pgaptmp_000409 | pgaptmp_009758 | pgaptmp_009919 | pgaptmp_009937 |
| hypothetical protein | HrpE/YscL family type III secretion apparatus protein | pgaptmp_009795 | pgaptmp_000411 | pgaptmp_009760 | pgaptmp_009921 | pgaptmp_009939 |
| type III secretion inner membrane ring lipoprotein SctJ | - | pgaptmp_009797 | pgaptmp_000413 | pgaptmp_009762 | pgaptmp_009923 | pgaptmp_009941 |
| EscI/YscI/HrpB family type III secretion system inner rod protein | - | pgaptmp_009798 | pgaptmp_000414 | pgaptmp_009763 | pgaptmp_009924 | pgaptmp_009942 |
| IS4 family transposase | - | - | pgaptmp_000419 | - | - | - |
| hypothetical protein | type III effector protein | pgaptmp_009805 | - | pgaptmp_009770 | pgaptmp_009931 | pgaptmp_009949 |

| | | | | | | |
|---|---|---|---|---|---|---|
| SycD/LcrH family type III secretion system chaperone | - | pgaptmp_009806 | - | pgaptmp_009771 | pgaptmp_009932 | pgaptmp_009950 |
| hypothetical protein | type III effector protein | pgaptmp_009807 | - | pgaptmp_009772 | pgaptmp_009933 | pgaptmp_009951 |
| hypothetical protein | type III effector protein | pgaptmp_009808 | - | pgaptmp_009773 | pgaptmp_009934 | pgaptmp_009952 |
| hypothetical protein | type III effector protein | pgaptmp_009809 | - | pgaptmp_009774 | pgaptmp_009935 | pgaptmp_009953 |
| hypothetical protein | gspD | pgaptmp_009810 | - | pgaptmp_009775 | pgaptmp_009936 | pgaptmp_009954 |
| recombinase family protein | - | pgaptmp_009814 | pgaptmp_000421 | pgaptmp_009779 | pgaptmp_009940 | pgaptmp_009958 |
| hypothetical protein | Effector protein - pseudomonas | pgaptmp_009817 | pgaptmp_000424 | pgaptmp_009782 | pgaptmp_009943 | pgaptmp_009962 |
| hypothetical protein | type II toxin-antitoxin system Phd/YefM family antitoxin | pgaptmp_009832 | pgaptmp_000439 | pgaptmp_009797 | pgaptmp_009958 | pgaptmp_009977 |
| type II toxin-antitoxin system prevent-host-death family antitoxin | - | pgaptmp_009833 | pgaptmp_000440 | pgaptmp_009798 | pgaptmp_009959 | pgaptmp_009978 |
| type II toxin-antitoxin system VapC family toxin | - | pgaptmp_009679 | pgaptmp_000295 | pgaptmp_009644 | pgaptmp_009803 | pgaptmp_009823 |
| hypothetical protein | type III effector protein | pgaptmp_009738 | pgaptmp_000354 | pgaptmp_009703 | pgaptmp_009862 | pgaptmp_009881 |
| transposase | - | pgaptmp_009756 | pgaptmp_000372 | pgaptmp_009721 | pgaptmp_009880 | pgaptmp_009899 |
| type III secretion system export apparatus subunit SctV | - | pgaptmp_009762 | pgaptmp_000378 | pgaptmp_009727 | pgaptmp_009886 | pgaptmp_009905 |
| type III secretion system ATPase SctN | - | pgaptmp_009794 | pgaptmp_000410 | pgaptmp_009759 | pgaptmp_009920 | pgaptmp_009938 |
| **Annotation** | **Conserved domain/protein** | **pBjCNPSo22b** | **pBjCNPSo29b** | **pBjCNPSo31b** | **pBjCNPSo34b** | **pBjCNPSo38b** |

| | | | | | | |
|---|---|---|---|---|---|---|
| type II toxin-antitoxin system VapC family toxin | - | pgaptmp_009389 | pgaptmp_000004 | pgaptmp_009355 | pgaptmp_000004 | pgaptmp_009522 |
| type II toxin-antitoxin system VapB family antitoxin | - | pgaptmp_009390 | pgaptmp_000005 | pgaptmp_009356 | pgaptmp_000005 | pgaptmp_009523 |
| type II toxin-antitoxin system VapC family toxin | - | pgaptmp_009392 | pgaptmp_000007 | pgaptmp_009358 | pgaptmp_000007 | pgaptmp_009525 |
| type II toxin-antitoxin system Phd/YefM family antitoxin | - | pgaptmp_009393 | pgaptmp_000008 | pgaptmp_009359 | pgaptmp_000008 | pgaptmp_009526 |
| hypothetical protein | non-ribosomal peptide synthetase family protein | pgaptmp_009425 | pgaptmp_000040 | pgaptmp_009392 | pgaptmp_000040 | pgaptmp_009558 |
| C48 family peptidase | | pgaptmp_009430 | - | pgaptmp_009397 | - | pgaptmp_009564 |
| hypothetical protein | E3 ubiquitin transferase SlrP | pgaptmp_009433 | pgaptmp_000049 | pgaptmp_009400 | pgaptmp_000049 | pgaptmp_009567 |
| hypothetical protein | HlyD family efflux transporter | pgaptmp_009435 | pgaptmp_000051 | pgaptmp_009402 | pgaptmp_000051 | pgaptmp_009569 |
| cold-shock protein | CspA family | pgaptmp_009438 | pgaptmp_000054 | pgaptmp_009405 | pgaptmp_000054 | pgaptmp_009572 |
| pyridoxal-phosphate dependent enzyme | cysK | pgaptmp_009441 | pgaptmp_000057 | pgaptmp_009408 | pgaptmp_000057 | pgaptmp_009575 |
| nucleotidyl transferase AbiEii/AbiGii toxin family protein | - | pgaptmp_009451 | pgaptmp_000067 | pgaptmp_009418 | pgaptmp_000067 | pgaptmp_009585 |
| GNAT family N-acetyltransferase | - | pgaptmp_009471 | pgaptmp_000087 | pgaptmp_009437 | pgaptmp_000086 | - |
| hypothetical protein | class I SAM-dependent methyltransferase | pgaptmp_009475 | pgaptmp_000091 | pgaptmp_009441 | pgaptmp_000090 | pgaptmp_009609 |
| protein-L-isoaspartate(D-aspartate) O-methyltransferase | - | pgaptmp_009476 | pgaptmp_000092 | pgaptmp_009442 | pgaptmp_000091 | pgaptmp_009610 |

| | | | | | | |
|---|---|---|---|---|---|---|
| N-formylglutamate amidohydrolase | formylglutamate hydrolase (FGase) | pgaptmp_009477 | pgaptmp_000093 | pgaptmp_009443 | pgaptmp_000092 | pgaptmp_009611 |
| chaperonin GroEL | - | pgaptmp_009483 | pgaptmp_000099 | pgaptmp_009449 | pgaptmp_000098 | pgaptmp_009617 |
| co-chaperone GroES | - | pgaptmp_009484 | pgaptmp_000100 | pgaptmp_009450 | pgaptmp_000099 | pgaptmp_009618 |
| nucleotidyl transferase AbiEii/AbiGii toxin family protein | - | pgaptmp_009503 | pgaptmp_000119 | pgaptmp_009469 | pgaptmp_000118 | pgaptmp_009637 |
| GMC family oxidoreductase | choline dehydrogenase BetA | pgaptmp_009516 | pgaptmp_000132 | pgaptmp_009482 | pgaptmp_000131 | pgaptmp_009650 |
| MFS transporter | - | pgaptmp_009518 | pgaptmp_000134 | pgaptmp_009484 | pgaptmp_000133 | pgaptmp_009652 |
| dihydroxy-acid dehydratase | - | pgaptmp_009525 | pgaptmp_000141 | pgaptmp_009491 | pgaptmp_000140 | pgaptmp_009659 |
| tripartite tricarboxylate transporter permease | - | pgaptmp_009530 | pgaptmp_000146 | pgaptmp_009496 | pgaptmp_000145 | pgaptmp_009664 |
| tripartite tricarboxylate transporter permease | - | pgaptmp_009531 | pgaptmp_000147 | pgaptmp_009497 | pgaptmp_000146 | pgaptmp_009665 |
| tripartite tricarboxylate transporter permease | - | pgaptmp_009532 | pgaptmp_000148 | pgaptmp_009498 | pgaptmp_000147 | pgaptmp_009666 |
| cystathionine gamma-synthase family protein | - | pgaptmp_009536 | pgaptmp_000152 | pgaptmp_009502 | pgaptmp_000151 | pgaptmp_009670 |
| E3 ubiquitin--protein ligase | - | pgaptmp_009537 | pgaptmp_000153 | pgaptmp_009503 | pgaptmp_000152 | pgaptmp_009671 |
| calcium-binding protein | - | pgaptmp_009567 | pgaptmp_000184 | pgaptmp_009532 | pgaptmp_000173 | pgaptmp_009702 |
| methyltransferase domain-containing protein | class I SAM-dependent methyltransferase | pgaptmp_009568 | pgaptmp_000185 | pgaptmp_009533 | pgaptmp_000174 | pgaptmp_009703 |
| peptidase domain-containing ABC transporter | HlyB | pgaptmp_009569 | - | pgaptmp_009534 | pgaptmp_000175 | pgaptmp_009704 |
| HlyD family type I secretion periplasmic adaptor subunit | - | pgaptmp_009570 | pgaptmp_000187 | pgaptmp_009535 | pgaptmp_000176 | pgaptmp_009705 |

| | | | | | | |
|---|---|---|---|---|---|---|
| glycoside hydrolase family 99-like domain-containing protein | - | pgaptmp_009572 | pgaptmp_000189 | pgaptmp_009537 | pgaptmp_000178 | pgaptmp_009707 |
| dTDP-glucose 4,6-dehydratase | - | pgaptmp_009577 | pgaptmp_000194 | pgaptmp_009542 | pgaptmp_000183 | pgaptmp_009712 |
| glucose-1-phosphate thymidylyltransferase RfbA | - | pgaptmp_009578 | pgaptmp_000195 | pgaptmp_009543 | pgaptmp_000184 | pgaptmp_009713 |
| dTDP-4-dehydrorhamnose 3,5-epimerase | - | pgaptmp_009579 | pgaptmp_000196 | pgaptmp_009544 | pgaptmp_000185 | pgaptmp_009714 |
| dTDP-4-dehydrorhamnose reductase | - | pgaptmp_009580 | pgaptmp_000197 | pgaptmp_009545 | pgaptmp_000186 | pgaptmp_009715 |
| glycosyltransferase family 2 protein | - | pgaptmp_009581 | pgaptmp_000198 | pgaptmp_009546 | pgaptmp_000187 | pgaptmp_009716 |
| glycosyltransferase family 2 protein | - | pgaptmp_009582 | pgaptmp_000199 | pgaptmp_009547 | pgaptmp_000188 | pgaptmp_009717 |
| glycosyltransferase family 4 protein | - | pgaptmp_009584 | pgaptmp_000201 | pgaptmp_009549 | pgaptmp_000190 | pgaptmp_009719 |
| glycosyltransferase family 4 protein | - | pgaptmp_009593 | pgaptmp_000210 | pgaptmp_009558 | pgaptmp_000199 | pgaptmp_009728 |
| ABC transporter permease | TagG | pgaptmp_009595 | pgaptmp_000212 | pgaptmp_009560 | pgaptmp_000201 | pgaptmp_009730 |
| ABC transporter ATP-binding protein | TagH | pgaptmp_009596 | pgaptmp_000213 | pgaptmp_009561 | pgaptmp_000202 | pgaptmp_009731 |
| glycosyltransferase | - | pgaptmp_009597 | pgaptmp_000214 | pgaptmp_009562 | pgaptmp_000203 | pgaptmp_009732 |
| glycosyltransferase | - | pgaptmp_009598 | pgaptmp_000215 | pgaptmp_009563 | pgaptmp_000204 | pgaptmp_009733 |
| glycosyltransferase | - | - | - | pgaptmp_009564 | - | - |
| acetyltransferase | sialic acid O-acetyltransferase NeuD family | pgaptmp_009599 | pgaptmp_000216 | pgaptmp_009565 | pgaptmp_000205 | pgaptmp_009734 |
| carotenoid biosynthesis protein | - | pgaptmp_009606 | pgaptmp_000223 | pgaptmp_009572 | pgaptmp_000212 | pgaptmp_009741 |
| EAL domain-containing protein | response regulator (Signal | pgaptmp_009631 | pgaptmp_000247 | pgaptmp_009596 | pgaptmp_000236 | pgaptmp_009766 |

| | | | | | | |
|---|---|---|---|---|---|---|
| | transduction mechanisms) | | | | | |
| hypothetical protein | GNAT acetyltransferase | pgaptmp_009635 | pgaptmp_000251 | pgaptmp_009600 | pgaptmp_000240 | pgaptmp_009770 |
| EAL domain-containing protein | response regulator (Signal transduction mechanisms) | pgaptmp_009646 | pgaptmp_000262 | pgaptmp_009611 | pgaptmp_000251 | pgaptmp_009781 |
| GNAT family N-acetyltransferase | - | - | - | - | - | pgaptmp_009605 |
| Gfo/Idh/MocA family oxidoreductase | - | pgaptmp_009515 | pgaptmp_000131 | pgaptmp_009481 | pgaptmp_000130 | pgaptmp_009649 |
| peptidase domain-containing ABC transporter | - | - | pgaptmp_000186 | - | - | - |
| DegT/DnrJ/EryC1/StrS family aminotransferase | - | pgaptmp_009594 | pgaptmp_000211 | pgaptmp_009559 | pgaptmp_000200 | pgaptmp_009729 |
| LysR substrate-binding domain-containing protein | - | pgaptmp_009615 | pgaptmp_000232 | pgaptmp_009581 | pgaptmp_000221 | pgaptmp_009750 |
| GGDEF domain-containing protein | - | pgaptmp_009644 | pgaptmp_000260 | pgaptmp_009609 | pgaptmp_000249 | pgaptmp_009779 |

**Table S2.** Accessory and unique genome of symbiosis island A of the *B. japonicum* group.

| **Accessory genome of the *B. japonicum* group** | | | | | | | | | |
|---|---|---|---|---|---|---|---|---|---|
| **Annotation** | **Strains** | | | | | | | | |
| | **CNPSo 17** | **CNPSo 22** | **CNPSo 23** | **CNPSo 24** | **CNPSo 29** | **CNPSo 31** | **CNPSo 34** | **CNPSo 38** | **CPAC 15** |
| IS21-like element ISFK1 family helper ATPase IstB | 17final_07730 | 22final_08005 | 23final_07707 | 24final_07699 | 29final_08460 | 31final_07983 | - | 38final_08138 | cpac15ori_final_07695 |
| IS21-like element ISFK1 family helper ATPase IstB | 17final_07917 | 22final_08190 | - | 24final_07886 | 29final_08643 | 31final_08168 | 34final_08604 | 38final_08324 | cpac15ori_final_07874 |
| putative transposase | 17final_07548 | 22final_07843 | 23final_07524 | 24final_07516 | 29final_08297 | 31final_07820 | 34final_08255 | 38final_07972 | - |
| IS3 family transposase ISOba1 | 17final_07435 | 22final_07728 | 23final_07411 | 24final_07403 | 29final_08182 | 31final_07703 | 34final_08143 | 38final_07859 | - |
| IS21 family transposase ISPpu7 | 17final_07860 | 22final_08135 | 23final_07837 | 24final_07829 | 29final_08588 | 31final_08113 | 34final_08547 | 38final_08268 | - |
| IS5 family transposase ISBj2 | 17final_07544 | 22final_07839 | 23final_07520 | 24final_07512 | 29final_08293 | 31final_07816 | 34final_08251 | 38final_07968 | - |
| IS5 family transposase ISBj2 | 17final_07547 | 22final_07842 | 23final_07523 | 24final_07515 | 29final_08296 | 31final_07819 | 34final_08254 | 38final_07971 | - |
| IS5 family transposase ISBj2 | 17final_07871 | 22final_08146 | 23final_07848 | 24final_07840 | 29final_08599 | 31final_08124 | 34final_08558 | 38final_08279 | - |
| IS3 family transposase ISSme1 | 17final_07434 | 22final_07727 | 23final_07410 | 24final_07402 | 29final_08181 | 31final_07702 | 34final_08142 | 38final_07858 | - |
| IS6 family transposase ISBj7 | 17final_07877 | 22final_08152 | 23final_07854 | 24final_07846 | 29final_08605 | 31final_08130 | 34final_08564 | 38final_08285 | - |
| IS6 family transposase ISBj7 | 17final_07885 | 22final_08159 | 23final_07862 | 24final_07854 | 29final_08612 | 31final_08137 | 34final_08572 | 38final_08292 | - |
| putative transposase | 17final_07876 | 22final_08151 | 23final_07853 | 24final_07845 | 29final_08604 | 31final_08129 | 34final_08563 | 38final_08284 | - |
| noeE-like protein - sulphotransferase | - | 22final_07766 | 23final_07449 | 24final_07441 | 29final_08221 | 31final_07741 | 34final_08181 | 38final_07897 | cpac15ori_final_07439 |
| hypothetical protein | - | 22final_07828 | 23final_07509 | 24final_07501 | 29final_08282 | 31final_07805 | 34final_08240 | 38final_07957 | cpac15ori_final_07499 |
| hypothetical protein | - | 22final_07829 | 23final_07510 | 24final_07502 | 29final_08283 | 31final_07806 | 34final_08241 | 38final_07958 | cpac15ori_final_07500 |

| | | | | | | | | | |
|---|---|---|---|---|---|---|---|---|---|
| hypothetical protein | 17final_07676 | 22final_07950 | - | 24final_07645 | 29final_08404 | 31final_07928 | 34final_08361 | 38final_08081 | cpac15ori_final_07644 |
| hypothetical protein | 17final_07843 | 22final_08118 | 23final_07820 | 24final_07812 | 29final_08573 | 31final_08096 | 34final_08530 | 38final_08251 | - |
| ribbon-helix-helix domains of transcription repressor CopG, nickel responsive transcription factor NikR | 17final_07728 | 22final_08003 | 23final_07705 | 24final_07697 | 29final_08458 | 31final_07981 | 34final_08414 | 38final_08136 | - |
| IS630 family transposase ISRj1 | 17final_07754 | 22final_08029 | 23final_07731 | 24final_07723 | 29final_08484 | 31final_08007 | 34final_08439 | 38final_08162 | - |
| IS21 family transposase ISPpu7 | 17final_07731 | 22final_08006 | 23final_07708 | 24final_07700 | 29final_08461 | 31final_07984 | - | 38final_08139 | - |
| hypothetical protein | - | 22final_08240 | 23final_07944 | - | 29final_08693 | 31final_08218 | 34final_08653 | 38final_08373 | cpac15ori_final_07923 |
| IS21-like element helper ATPase IstB | 17final_07840 | 22final_08115 | 23final_07817 | 24final_07809 | - | 31final_08093 | 34final_08527 | 38final_08248 | - |
| hypothetical protein | 17final_07527 | 22final_07821 | 23final_07503 | 24final_07495 | 29final_08275 | 31final_07798 | - | - | cpac15ori_final_07493 |
| hypothetical protein | 17final_07528 | 22final_07822 | 23final_07504 | 24final_07496 | 29final_08276 | 31final_07799 | - | - | cpac15ori_final_07494 |
| hypothetical protein | 17final_07844 | 22final_08119 | 23final_07821 | 24final_07813 | - | 31final_08097 | 34final_08531 | 38final_08252 | - |
| hypothetical protein | 17final_07845 | 22final_08120 | 23final_07822 | 24final_07814 | - | 31final_08098 | 34final_08532 | 38final_08253 | - |
| hypothetical protein | 17final_07846 | 22final_08121 | 23final_07823 | 24final_07815 | - | 31final_08099 | 34final_08533 | 38final_08254 | - |
| IS3 family transposase ISRj2 | 17final_07815 | 22final_08090 | 23final_07792 | 24final_07784 | 29final_08545 | 31final_08068 | - | - | - |
| putative transposase | 17final_07816 | 22final_08091 | 23final_07793 | 24final_07785 | 29final_08546 | 31final_08069 | - | - | - |
| putative transposase | - | 22final_07996 | 23final_07697 | - | 29final_08451 | 31final_07974 | 34final_08407 | 38final_08129 | - |
| hypothetical protein | 17final_07738 | - | 23final_07715 | 24final_07707 | - | - | 34final_08423 | 38final_08146 | cpac15ori_final_07703 |
| hypothetical protein | 17final_07531 | 22final_07825 | - | - | 29final_08279 | 31final_07802 | 34final_08237 | 38final_07954 | - |

| | | | | | | | | | |
|---|---|---|---|---|---|---|---|---|---|
| putative transposase | - | 22final_07881 | - | - | 29final_08335 | 31final_07858 | 34final_08293 | 38final_08011 | - |
| group II intron reverse transcriptase/maturase | - | 22final_07880 | - | - | 29final_08334 | 31final_07857 | 34final_08292 | 38final_08010 | - |
| IS5 family transposase ISBj2 | - | 22final_07882 | - | - | 29final_08336 | 31final_07859 | 34final_08294 | 38final_08012 | - |
| Protein of unknown function (DUF3551) | - | 22final_07737 | - | - | 29final_08192 | 31final_07712 | 34final_08152 | 38final_07868 | - |
| hypothetical protein | - | 22final_07820 | - | - | 29final_08274 | 31final_07797 | 34final_08234 | 38final_07951 | - |
| putative transposase | - | 22final_07883 | - | - | 29final_08337 | 31final_07860 | 34final_08295 | 38final_08013 | - |
| reverse transcriptase family protein | - | 22final_07884 | - | - | 29final_08338 | 31final_07861 | 34final_08296 | 38final_08014 | - |
| hypothetical protein | - | 22final_07885 | - | - | 29final_08339 | 31final_07863 | 34final_08297 | 38final_08015 | - |
| hypothetical protein | - | 22final_07887 | - | - | 29final_08341 | 31final_07865 | 34final_08299 | 38final_08017 | - |
| hypothetical protein | - | 22final_07888 | - | - | 29final_08342 | 31final_07866 | 34final_08300 | 38final_08018 | - |
| Purine efflux pump PbuE/Predicted arabinose efflux permease, MFS family [Carbohydrate transport and metabolism] | - | 22final_07889 | - | - | 29final_08343 | 31final_07867 | 34final_08301 | 38final_08019 | - |
| IS110 family transposase ISPye17 | - | 22final_07890 | - | - | 29final_08344 | 31final_07868 | 34final_08302 | 38final_08020 | - |
| putative transposase | - | 22final_07891 | - | - | 29final_08345 | 31final_07869 | 34final_08303 | 38final_08021 | - |
| Cold shock protein CspA | - | 22final_07892 | - | - | 29final_08346 | 31final_07870 | 34final_08304 | 38final_08022 | - |
| prolyl oligopeptidase family protein (preP) | - | 22final_07893 | - | - | 29final_08347 | 31final_07871 | 34final_08305 | 38final_08023 | - |
| C-terminal DNA-binding domain of LuxR-like proteins | - | 22final_07894 | - | - | 29final_08348 | 31final_07872 | 34final_08306 | 38final_08024 | - |
| Phosphonopyruvate hydrolase | - | 22final_07895 | - | - | 29final_08349 | 31final_07873 | 34final_08307 | 38final_08025 | - |

| | | | | | | | | | |
|---|---|---|---|---|---|---|---|---|---|
| Phosphocholine cytidylyltransferases | - | 22final_07896 | - | - | 29final_08350 | 31final_07874 | 34final_08308 | 38final_08026 | - |
| phosphonates metabolism transcriptional regulator PhnF | - | 22final_07897 | - | - | 29final_08351 | 31final_07875 | 34final_08309 | 38final_08027 | - |
| Type I phosphodiesterase / nucleotide pyrophosphatase | - | 22final_07898 | - | - | 29final_08352 | 31final_07876 | 34final_08310 | 38final_08028 | - |
| ABC-type nickel/oligopeptide-like import system | - | 22final_07899 | - | - | 29final_08353 | 31final_07877 | 34final_08311 | 38final_08029 | - |
| ABC-type dipeptide/oligopeptide/nickel transport system | - | 22final_07900 | - | - | 29final_08354 | 31final_07878 | 34final_08312 | 38final_08030 | - |
| sugar isomerase domain-containing protein | - | 22final_07912 | - | - | 29final_08366 | 31final_07890 | 34final_08324 | 38final_08044 | - |
| hypothetical protein | - | 22final_07992 | - | - | 29final_08447 | 31final_07970 | 34final_08403 | 38final_08125 | - |
| peptidase M16 family protein | - | 22final_08041 | - | - | 29final_08496 | 31final_08019 | 34final_08451 | 38final_08174 | - |
| hypothetical protein | - | 22final_07988 | - | - | 29final_08443 | 31final_07966 | 34final_08399 | 38final_08121 | - |
| putative type III system effector | 17final_07714 | - | 23final_07690 | 24final_07683 | - | - | - | 38final_08119 | cpac15ori_final_07682 |
| IS5 family transposase ISAcp3 | - | 22final_07886 | - | - | 29final_08340 | 31final_07864 | 34final_08298 | 38final_08016 | - |
| recombinase family protein | - | 22final_08002 | - | - | 29final_08457 | 31final_07980 | 34final_08413 | 38final_08135 | - |
| hypothetical protein | 17final_07388 | - | - | - | 29final_08138 | 31final_07659 | 34final_08099 | 38final_07815 | - |
| group II intron reverse transcriptase/maturase | 17final_07417 | - | 23final_07393 | 24final_07385 | - | - | - | - | cpac15ori_final_07384 |
| putative transposase | 17final_07431 | - | 23final_07407 | 24final_07399 | - | - | - | - | cpac15ori_final_07398 |
| putative transposase | 17final_07884 | - | 23final_07861 | 24final_07853 | - | - | 34final_08571 | - | - |
| D-alanine--D-alanyl carrier protein ligase | 17final_07447 | - | 23final_07423 | 24final_07415 | - | - | - | - | cpac15ori_final_07413 |

| | | | | | | | | | |
|---|---|---|---|---|---|---|---|---|---|
| hypothetical protein | 17final_07893 | - | 23final_07870 | 24final_07862 | - | - | - | - | cpac15ori_final_07850 |
| radical SAM family RiPP maturation amino acid epimerase | 17final_07633 | - | 23final_07610 | 24final_07602 | - | - | - | - | cpac15ori_final_07601 |
| hypothetical protein | 17final_07526 | - | 23final_07502 | 24final_07494 | - | - | - | - | cpac15ori_final_07492 |
| Alpha-ketoglutarate-dependent sulfate ester dioxygenase | 17final_07585 | - | 23final_07561 | 24final_07553 | - | - | - | - | cpac15ori_final_07551 |
| Oxaloacetate decarboxylase | 17final_07595 | - | 23final_07571 | 24final_07563 | - | - | - | - | cpac15ori_final_07561 |
| Glycine cleavage system transcriptional activator/LysR family transcriptional regulator | 17final_07596 | - | 23final_07572 | 24final_07564 | - | - | - | - | cpac15ori_final_07562 |
| hypothetical protein | 17final_07597 | - | 23final_07573 | 24final_07565 | - | - | - | - | cpac15ori_final_07563 |
| helix-turn-helix transcriptional regulator | 17final_07598 | - | 23final_07574 | 24final_07566 | - | - | - | - | cpac15ori_final_07564 |
| hypothetical protein | 17final_07599 | - | 23final_07575 | 24final_07567 | - | - | - | - | cpac15ori_final_07565 |
| putative transposase | 17final_07600 | - | 23final_07576 | 24final_07568 | - | - | - | - | cpac15ori_final_07566 |
| Alpha-ketoglutarate-dependent sulfate ester dioxygenase | 17final_07601 | - | 23final_07577 | 24final_07569 | - | - | - | - | cpac15ori_final_07567 |
| hypothetical protein | 17final_07604 | - | 23final_07580 | 24final_07572 | - | - | - | - | cpac15ori_final_07570 |
| hypothetical protein | 17final_07605 | - | 23final_07581 | 24final_07573 | - | - | - | - | cpac15ori_final_07571 |
| hypothetical protein | 17final_07606 | - | 23final_07582 | 24final_07574 | - | - | - | - | cpac15ori_final_07572 |
| hypothetical protein | 17final_07607 | - | 23final_07583 | 24final_07575 | - | - | - | - | cpac15ori_final_07573 |
| hypothetical protein | 17final_07608 | - | 23final_07584 | 24final_07576 | - | - | - | - | cpac15ori_final_07574 |
| nucleotidyl transferase AbiEii/AbiGii toxin family protein | 17final_07609 | - | 23final_07585 | 24final_07577 | - | - | - | - | cpac15ori_final_07575 |

| | | | | | | | | | |
|---|---|---|---|---|---|---|---|---|---|
| nucleotidyl transferase AbiEii/AbiGii toxin family protein | 17final_07610 | - | 23final_07586 | 24final_07578 | - | - | - | - | cpac15ori_final_07576 |
| type IV toxin-antitoxin system AbiEi family antitoxin domain-containing protein | 17final_07611 | - | 23final_07587 | 24final_07579 | - | - | - | - | cpac15ori_final_07577 |
| nucleotidyl transferase AbiEii/AbiGii toxin family protein | 17final_07612 | - | 23final_07588 | 24final_07580 | - | - | - | - | cpac15ori_final_07578 |
| hypothetical protein | 17final_07613 | - | 23final_07589 | 24final_07581 | - | - | - | - | cpac15ori_final_07579 |
| hypothetical protein | 17final_07614 | - | 23final_07590 | 24final_07582 | - | - | - | - | cpac15ori_final_07580 |
| hypothetical protein | 17final_07615 | - | 23final_07591 | 24final_07583 | - | - | - | - | cpac15ori_final_07581 |
| KilA-N domain containing protein | 17final_07616 | - | 23final_07592 | 24final_07584 | | | | | cpac15ori_final_07582 |
| hypothetical protein | 17final_07617 | - | 23final_07593 | 24final_07585 | | | | | cpac15ori_final_07583 |
| hypothetical protein | 17final_07618 | - | 23final_07594 | 24final_07586 | | | | | cpac15ori_final_07584 |
| IS1595 family transposase ISNwi1 | 17final_07619 | - | 23final_07595 | 24final_07587 | | | | | cpac15ori_final_07585 |
| hypothetical protein | 17final_07620 | - | 23final_07596 | 24final_07588 | | | | | cpac15ori_final_07586 |
| hypothetical protein | 17final_07621 | - | 23final_07597 | 24final_07589 | | | | | cpac15ori_final_07587 |
| Lrp/AsnC family transcriptional regulator | 17final_07622 | - | 23final_07598 | 24final_07590 | | | | | cpac15ori_final_07588 |
| FAD-dependent oxidoreductase | 17final_07623 | - | 23final_07599 | 24final_07591 | | | | | cpac15ori_final_07589 |
| Phosphoenolpyruvate hydrolase-like | 17final_07624 | - | 23final_07600 | 24final_07592 | | | | | cpac15ori_final_07590 |
| hypothetical protein | 17final_07625 | - | 23final_07601 | 24final_07593 | | | | | cpac15ori_final_07591 |
| hypothetical protein | 17final_07626 | - | 23final_07602 | 24final_07594 | | | | | cpac15ori_final_07592 |

| | | | | | | | | | |
|---|---|---|---|---|---|---|---|---|---|
| NAD(P)/FAD-dependent oxidoreductase | 17final_07627 | - | 23final_07604 | 24final_07596 | | | | | cpac15ori_final_07595 |
| NAD(P)/FAD-dependent oxidoreductase | 17final_07628 | - | 23final_07605 | 24final_07597 | | | | | cpac15ori_final_07596 |
| hypothetical protein | 17final_07629 | - | 23final_07606 | 24final_07598 | | | | | cpac15ori_final_07597 |
| hypothetical protein | 17final_07634 | - | 23final_07611 | 24final_07603 | | | | | cpac15ori_final_07602 |
| hypothetical protein | 17final_07635 | - | 23final_07612 | 24final_07604 | | | | | cpac15ori_final_07603 |
| hypothetical protein | 17final_07636 | - | 23final_07613 | 24final_07605 | | | | | cpac15ori_final_07604 |
| hypothetical protein | 17final_07637 | - | 23final_07614 | 24final_07606 | | | | | cpac15ori_final_07605 |
| hypothetical protein | 17final_07638 | | 23final_07615 | 24final_07607 | | | | | cpac15ori_final_07606 |
| Transcriptional regulator HipB | 17final_07639 | - | 23final_07616 | 24final_07608 | - | - | - | - | cpac15ori_final_07607 |
| type II toxin-antitoxin sytem toxin HipA | 17final_07640 | - | 23final_07617 | 24final_07609 | - | - | - | - | cpac15ori_final_07608 |
| hypothetical protein | 17final_07641 | - | 23final_07618 | 24final_07610 | - | - | - | - | cpac15ori_final_07609 |
| hypothetical protein | 17final_07642 | - | 23final_07619 | 24final_07611 | - | - | - | - | cpac15ori_final_07610 |
| hypothetical protein | 17final_07766 | - | 23final_07743 | 24final_07735 | - | - | - | - | cpac15ori_final_07731 |
| IS630 family transposase ISRj1 | - | 22final_07929 | - | - | 29final_08383 | 31final_07907 | - | - | - |
| IS3 family transposase ISRj2 | 17final_07432 | - | - | 24final_07400 | - | - | - | - | cpac15ori_final_07399 |
| IS3 family transposase ISRj2 | - | - | - | - | - | - | 34final_08502 | 38final_08223 | cpac15ori_final_07780 |
| hypothetical protein | - | 22final_07778 | - | - | - | 31final_07754 | - | 38final_07909 | - |
| hypothetical protein | - | 22final_08013 | - | - | 29final_08468 | 31final_07991 | - | - | - |

| | | | | | | | | | |
|---|---|---|---|---|---|---|---|---|---|
| hypothetical protein | - | 22final_08203 | - | - | 29final_08656 | 31final_08181 | - | - | - |
| hypothetical protein | - | - | 23final_07603 | 24final_07595 | - | - | - | - | cpac15ori_final_07594 |
| hypothetical protein | 17final_07726 | - | 23final_07703 | 24final_07695 | - | - | - | - | - |
| hypothetical protein | 17final_07727 | - | 23final_07704 | 24final_07696 | - | - | - | - | - |
| putative transposase | - | - | - | - | - | - | 34final_08503 | 38final_08224 | - |
| putative transposase | - | - | - | - | - | - | 34final_08586 | 38final_08306 | - |
| Ulp1 family isopeptidase/ putative *nopD2* | - | - | - | - | 29final_08574 | - | - | - | cpac15ori_final_07807 |
| | **Unique genome of the *B. japonicum* group** | | | | | | | | |
| **Annotation** | **Strains** | | | | | | | | |
| | **CNPSo 17** | **CNPSo 22** | **CNPSo 23** | **CNPSo 24** | **CNPSo 29** | **CNPSo 31** | **CNPSo 34** | **CNPSo 38** | **CPAC 15** |
| IS3 family transposase ISRj2 | - | - | - | - | - | - | 34final_08483 | - | - |
| putative transposase | - | - | - | - | - | - | 34final_08484 | - | - |
| IS1380 family transposase ISBdi2 | - | - | - | - | - | - | - | 38final_08042 | - |
| IS1380 family transposase ISBdi2 | - | - | - | - | - | - | - | 38final_08120 | - |
| IS1380 family transposase ISBdi2 | - | - | - | - | - | 31final_07745 | - | - | - |
| hypothetical protein | - | - | 23final_07894 | - | - | - | - | - | - |
| hypothetical protein | - | - | 23final_07895 | - | - | - | - | - | - |
| LuxR C-terminal-related transcriptional regulator | - | - | - | - | - | 31final_07790 | - | - | - |

| | | | | | | | | | |
|---|---|---|---|---|---|---|---|---|---|
| putative transposase | - | - | - | - | - | 31final_07862 | - | - | - |
| IS21-like element helper ATPase IstB | - | - | - | - | 29final_08570 | - | - | - | - |
| putative transposase | - | - | - | - | - | - | - | - | cpac15ori_final_07510 |
| putative transposase | - | - | - | - | - | - | - | - | cpac15ori_final_07513 |
| putative transposase | - | - | - | - | - | - | - | - | cpac15ori_final_07514 |
| hypothetical protein | - | - | - | - | - | - | - | - | cpac15ori_final_07593 |
| hypothetical protein | - | - | - | - | 29final_08189 | - | - | - | - |
| putative E3 ubiquitin-protein ligase ipaH7.8/ putative *nopM* | - | - | - | - | - | - | - | 38final_07999 | - |
| IS256 family transposase ISMex3 | - | - | - | - | - | - | - | - | - |
| IS3 family transposase ISRj2 | - | - | - | - | - | - | - | - | cpac15ori_final_07781 |
| IS3 family transposase ISRj2 | - | - | 23final_07408 | - | - | - | - | - | - |
| hypothetical protein | 17final_07473 | - | - | - | - | - | - | - | - |
| ATPase domain | 17final_07534 | - | - | - | - | - | - | - | - |
| IS6 family transposase ISBj7 | - | - | - | - | - | - | - | - | cpac15ori_final_07836 |
| hypothetical protein | - | - | - | - | - | - | - | 38final_08043 | - |

**Table S3.** Accessory and unique genome of symbiosis island A of the *B. diazoefficiens* group.

| | **Accessory genome of the *B. diazoefficiens* group** | | | | | | | | |
|---|---|---|---|---|---|---|---|---|---|
| **Annotation** | **Strains** | | | | | | | | |
| | **CNPSo 10** | **CNPSo 104** | **CNPSo 105** | **CNPSo 106** | **CNPSo 107** | **CNPSo 108** | **CNPSo 109** | **CNPSo 110** | **CPAC 7** |
| hypothetical protein | 10final_ 07144 | 104final_ 07144 | 105final_ 07141 | 106final_ 07151 | 107final_ 07145 | 108final_ 07139 | 109final_ 07143 | - | cpac7ori_fin al_07143 |
| hypothetical protein | 10final_ 07254 | 104final_ 07254 | 105final_ 07251 | - | 107final_ 07255 | 108final_ 07250 | 109final_ 07254 | 110final_ 07249 | cpac7ori_fin al_07255 |
| IS630 family transposase ISRj1 | 10final_ 07255 | 104final_ 07255 | 105final_ 07252 | - | 107final_ 07256 | 108final_ 07251 | 109final_ 07255 | 110final_ 07250 | cpac7ori_fin al_07256 |
| hypothetical protein | 10final_ 07256 | 104final_ 07256 | 105final_ 07253 | - | 107final_ 07257 | 108final_ 07252 | 109final_ 07256 | 110final_ 07251 | cpac7ori_fin al_07257 |
| hypothetical protein | 10final_ 07257 | 104final_ 07257 | 105final_ 07254 | - | 107final_ 07258 | 108final_ 07253 | 109final_ 07257 | 110final_ 07252 | cpac7ori_fin al_07258 |
| Acetyltransferase (GNAT) domain | 10final_ 07258 | 104final_ 07258 | 105final_ 07255 | - | 107final_ 07259 | 108final_ 07254 | 109final_ 07258 | 110final_ 07253 | cpac7ori_fin al_07259 |
| E3 ubiquitin-protein ligase ipaH3/putative *nopM* | 10final_ 07259 | 104final_ 07259 | 105final_ 07256 | - | 107final_ 07260 | 108final_ 07255 | 109final_ 07259 | 110final_ 07254 | cpac7ori_fin al_07260 |
| SbmA/BacA-like family | 10final_ 07260 | 104final_ 07260 | 105final_ 07257 | - | 107final_ 07261 | 108final_ 07256 | 109final_ 07260 | 110final_ 07255 | cpac7ori_fin al_07261 |
| Peptide antibiotic transporter SbmA | 10final_ 07261 | 104final_ 07261 | 105final_ 07258 | - | 107final_ 07262 | 108final_ 07257 | 109final_ 07261 | 110final_ 07256 | cpac7ori_fin al_07262 |
| SbmA/BacA-like family | 10final_ 07262 | 104final_ 07262 | 105final_ 07259 | - | 107final_ 07263 | 108final_ 07258 | 109final_ 07262 | 110final_ 07257 | cpac7ori_fin al_07263 |
| hypothetical protein | 10final_ 07265 | 104final_ 07265 | 105final_ 07262 | - | 107final_ 07266 | 108final_ 07261 | 109final_ 07265 | 110final_ 07260 | cpac7ori_fin al_07266 |
| IS3 family transposase ISAtu4 | 10final_ 07266 | 104final_ 07266 | 105final_ 07263 | - | 107final_ 07267 | 108final_ 07262 | 109final_ 07266 | 110final_ 07261 | cpac7ori_fin al_07267 |
| IS3 family transposase ISAtu4 | 10final_ 07267 | 104final_ 07267 | 105final_ 07264 | - | 107final_ 07268 | 108final_ 07263 | 109final_ 07267 | 110final_ 07262 | cpac7ori_fin al_07268 |
| IS110 family transposase ISWpi13 | 10final_ 07268 | 104final_ 07268 | 105final_ 07265 | - | 107final_ 07269 | 108final_ 07264 | 109final_ 07268 | 110final_ 07263 | cpac7ori_fin al_07269 |
| hypothetical protein | 10final_ 07269 | 104final_ 07269 | 105final_ 07266 | - | 107final_ 07270 | 108final_ 07265 | 109final_ 07269 | 110final_ 07264 | cpac7ori_fin al_07270 |

| | | | | | | | | | |
|---|---|---|---|---|---|---|---|---|---|
| LomR superfamily/outer membrane beta-barrel protein | 10final_07270 | 104final_07270 | 105final_07267 | - | 107final_07271 | 108final_07266 | 109final_07270 | 110final_07265 | cpac7ori_final_07271 |
| IS701 family transposase ISNha2 | 10final_07272 | 104final_07272 | 105final_07269 | - | 107final_07273 | 108final_07268 | 109final_07272 | 110final_07267 | cpac7ori_final_07273 |
| putative transposase | 10final_07349 | 104final_07350 | 105final_07346 | 106final_07336 | 107final_07350 | 108final_07345 | 109final_07349 | - | cpac7ori_final_07350 |
| putative transposase | 10final_07273 | 104final_07273 | 105final_07270 | - | 107final_07274 | 108final_07269 | 109final_07273 | 110final_07268 | cpac7ori_final_07274 |
| hypothetical protein | 10final_07487 | - | 105final_07484 | 106final_07474 | 107final_07488 | 108final_07483 | 109final_07487 | 110final_07481 | cpac7ori_final_07489 |
| hypothetical protein | 10final_07521 | 104final_07521 | 105final_07517 | 106final_07507 | 107final_07521 | - | 109final_07520 | 110final_07515 | cpac7ori_final_07522 |
| ISNCY family transposase ISBj12 | 10final_07474 | - | 105final_07471 | 106final_07461 | 107final_07475 | 108final_07470 | 109final_07474 | 110final_07468 | cpac7ori_final_07476 |
| hypothetical protein | 10final_07271 | 104final_07271 | 105final_07268 | - | 107final_07272 | 108final_07267 | 109final_07271 | 110final_07266 | cpac7ori_final_07272 |
| putative oxidoreductase | 10final_07086 | 104final_07086 | 105final_07083 | 106final_07093 | 107final_07087 | - | 109final_07085 | 110final_07080 | cpac7ori_final_07086 |
| hypothetical protein | 10final_07253 | 104final_07253 | 105final_07250 | - | 107final_07254 | 108final_07249 | 109final_07253 | - | cpac7ori_final_07254 |
| hypothetical protein | 10final_07520 | 104final_07520 | 105final_07516 | 106final_07506 | 107final_07520 | - | 109final_07519 | - | cpac7ori_final_07521 |
| hypothetical protein | 10final_07263 | - | 105final_07260 | - | 107final_07264 | 108final_07259 | 109final_07263 | 110final_07258 | cpac7ori_final_07264 |
| ISNCY family transposase ISBj12 | 10final_07264 | - | 105final_07261 | - | 107final_07265 | 108final_07260 | 109final_07264 | 110final_07259 | cpac7ori_final_07265 |
| group II intron reverse transcriptase/maturase | 10final_07125 | 104final_07125 | 105final_07122 | - | 107final_07126 | - | 109final_07124 | 110final_07119 | - |
| hypothetical protein | 10final_07312 | - | 105final_07309 | - | 107final_07313 | 108final_07308 | 109final_07312 | 110final_07307 | - |
| putative transposase | 10final_07186 | - | 105final_07183 | 106final_07194 | - | 108final_07182 | - | 110final_07180 | - |
| IS5 family transposase ISBj2 | - | 104final_07069 | - | 106final_07076 | - | 108final_07066 | 109final_07068 | - | cpac7ori_final_07069 |
| hypothetical protein | - | 104final_07186 | - | - | 107final_07187 | - | 109final_07186 | - | cpac7ori_final_07186 |

| | | | | | | | | | |
|---|---|---|---|---|---|---|---|---|---|
| sulfite exporter TauE/SafE family protein | - | - | - | 106final_07184 | - | - | 109final_07176 | 110final_07170 | cpac7ori_final_07176 |
| putative transposase | 10final_07069 | - | 105final_07066 | - | 107final_07070 | - | - | 110final_07063 | - |
| ISNCY family transposase ISBj12 | 10final_00656 | 104final_07475 | - | - | - | - | 109final_00658 | - | - |
| IS5 family transposase ISBj2 | - | 104final_07313 | - | 106final_07299 | - | - | - | - | cpac7ori_final_07313 |
| IS5 family transposase ISBj2 | - | - | - | - | - | 108final_07516 | - | 110final_07248 | - |
| | **Unique genome of the *B. diazoefficiens* group** | | | | | | | | |
| **Annotation** | **Strains** | | | | | | | | |
| | **CNPSo 10** | **CNPSo 104** | **CNPSo 105** | **CNPSo 106** | **CNPSo 107** | **CNPSo 108** | **CNPSo 109** | **CNPSo 110** | **CPAC 7** |
| IS1380 family transposase ISBdi2 | 10final_07504 | - | - | - | - | - | - | - | - |
| hypothetical protein | - | 104final_07263 | - | - | - | - | - | - | - |
| ISNCY family transposase ISBj12 | - | 104final_07264 | - | - | - | - | - | - | - |
| hypothetical protein | - | 104final_07301 | - | - | - | - | - | - | - |
| putative tranposase | - | 104final_07488 | - | - | - | - | - | - | - |
| IS1380 family transposase ISBdi2 | - | - | - | 106final_07099 | - | - | - | - | - |
| putative tranposase | - | - | - | - | - | 108final_07143 | - | - | - |
| putative tranposase | - | - | - | - | - | 108final_07515 | - | - | - |
| hypothetical protein | - | - | - | - | - | 108final_07555 | - | - | - |
| LPS-assembly protein (*LptD*) | - | - | - | - | - | - | - | 110final_07210 | - |

| | | | | | | | | | |
|---|---|---|---|---|---|---|---|---|---|
| putative tranposase | - | - | - | - | - | - | - | 110final_07513 | - |
| putative tranposase | - | - | - | - | - | - | - | 110final_07514 | - |
| IS21-like element helper ATPase IstB | - | - | - | - | - | - | - | - | cpac7ori_final_07227 |
| IS66 family transposase IS866 | - | - | - | - | - | - | - | - | cpac7ori_final_07461 |
